# *WRKY1* mediates transcriptional crosstalk between light and nitrogen signaling pathways in *Arabidopsis thaliana*

**DOI:** 10.1101/603142

**Authors:** Sachin Heerah, Manpreet Katari, Rebecca Penjor, Gloria Coruzzi, Amy Marshall-Colon

**Affiliations:** Department of Plant Biology, University of Illinois, 1201 W Gregory Dr, Urbana, IL 61801; Center for Genomics & Systems Biology, New York University, 12 Waverly Place, New York, NY 10001

## Abstract

Plant responses to multiple stimuli must be integrated to trigger transcriptional cascades that lead to changes in plant metabolism and development. Light (L) and nitrogen (N) are two signaling pathways that are intimately connected to each other and to plant energy status. Here, we describe the functional role of the WRKY1 transcription factor in mediating the regulation between L and N signaling pathways in *Arabidopsis thaliana*. WRKY1 participates in genome-wide transcriptional reprogramming in leaves in response to individual and combined L and N signals. A regulatory network was identified, consisting of 724 genes regulated by WRKY1 and involved in both N and L signaling pathways. The loss of WRKY1 gene function has marked effects on the L and N response of genes involved in N uptake and assimilation (primary metabolism) as well as stress response pathways (secondary metabolism). Our results support a model in which WRKY1 enables plants to activate genes involved in the recycling of cellular carbon resources when L is limiting but N is abundant, and up-regulate amino acid metabolism genes when both L and N are limiting. In this potential energy conservation mechanism, WRKY1 integrates responses to N and light-energy status to trigger changes in plant metabolism.

**Summary:** Based on transcriptome analysis, the *WRKY1* transcription factor mediates regulation of nitrogen and light signaling pathways in a potential energy conservation mechanism.

## INTRODUCTION

As sessile organisms, plants perceive multiple stimuli and must dynamically respond to complex environmental challenges in order to survive. Response to stimuli or stresses occur via signal transduction pathways that initiate transcriptional responses. Signal response pathways do not often act alone, but instead interact with other signaling pathways within a cell or tissue, ultimately resulting in emergent properties in the underlying gene regulatory networks. These pathways are likely connected via integrator molecules that mediate some common effects (Chen *et al*., 2013, Matiolli *et al*., 2011, Seo and Park, 2010). Light (L) and nitrogen (N) are two signaling pathways that are closely connected (Oliveira et al., 1999; Oliveira et al., 2001; Reed et al., 1983; Riens and Heldt, 1992; Chen et al., 2016). Nitrogen assimilation is dependent on reducing power and carbon skeletons derived from photosynthesis, while the photosynthetic apparatus is dependent on N availability to support the formation of chlorophyll and other components necessary for biomass accumulation (Blaesing *et al*., 2005; Lillo 2008; Matt et al., 2001a; Matt et al., 2001b). In addition, increasing evidence supports the notion that transcriptional crosstalk occurs between light and nitrogen signaling pathways in leaves to fine-tune plant energy status (Jonassen *et al*., 2008; Krouk *et al*., 2009; Nunes-Nesi *et al*., 2010; Obertello *et al*., 2010).

Knowledge about the regulation of genes commonly involved in L and N signaling pathways by transcription factors (TFs) is limited. Whole transcriptome analysis of Arabidopsis transcriptional response to different combinations of carbon (C), nitrogen, and light treatments revealed a change in expression of several genes, including a few known TFs (Krouk *et al*., 2009). That prior study revealed that 35% of the genome is controlled by L, C, or N signals or the combination of any of these signals (Krouk *et al*., 2009). A few shared elements in light and N signal transduction pathways have also been identified by studying the Arabidopsis bZIP TFs including bZIP1 (Obertello *et al*., 2010) and HY5/HYH (Jonassen *et al*., 2008). Genome-wide analysis of *bzip1* mutant seedlings revealed 33 genes with a significant interaction term for genotype, L, and N treatments, indicating that bZIP1 regulates a small group of genes involved in both L and N sensing (Obertello *et al*., 2010). HY5 and HYH were found to be essential for light-activated/phytochrome-mediated expression of nitrate reductase (Jonassen *et al*., 2008), in which the enhancement of NIA2 expression by light is dependent on HY5/HYH (Jonassen *et al*., 2009). It was also shown that HY5 is a shoot-to-root mobile TF that mediates the light activated N uptake by inducing expression of the NO_3_^−^ transporter NRT2.1 (Chen et al., 2016). Thus, despite the many and varied interactions between N and L signaling pathways, to our knowledge, only TFs from the bZIP gene family have been experimentally validated to integrate responses to both N and L signal transduction pathways.

During our previous work on Arabidopsis gene regulation by N status, we used network analysis to predict regulatory connections between genes and associated TFs (Gutierrez *et al*., 2008). We identified several TFs involved in positive and negative regulation of organic N metabolism and catabolism. Three regulatory hubs of an organic-N regulatory network identified were CCA1, GLK1, and bZIP1 (Gutierrez *et al*., 2008), each of which have been implicated in N and/or L signaling pathways (Wang and Tobin, 1998; Waters *et al*., 2009; Maekawa *et al*., 2015; Dietrich *et al*., 2011; Obertello *et al*., 2010). Indeed, independent experiments revealed that CCA1, GLK1 (Gutierrez *et al*., 2008), and bZIP1 (Gutierrez *et al*., 2008; Obertello *et al*., 2010; Baena-Gonzalez *et al*., 2008; Deitrich *et al*., 2011; Para *et al*., 2014) are involved in the regulation of genes in response to N and/or L signals. Our subsequent analysis of the N regulated genes described in Gutierrez *et al*. (2008) revealed an additional TF hub, WRKY1, which was also previously shown to be regulated by light (Krouk *et al*., 2009).

WRKY1 is a member of the WRKY family of TFs, which have diverse regulatory functions in response to biotic and abiotic stresses (Jia *et al*., 2015; Wei et al., 2008). WRKY TFs have been shown to activate or repress transcription and in some instances have dual activator/repressor functions. For example, OsWRKY72 and OsWRKY77 activate ABA signaling and repress GA signaling in rice (Xie *et al*., 2005). AtWRKY1 also plays a key role in the regulation of genes involved in ABA signaling and drought response in Arabidopsis (Qiao et al., 2016). Additional studies showed that other WRKY TFs respond to and also regulate gene response to light signals, where expression of AtWRKY22 is induced by light and repressed by dark treatment (Zhou *et al*., 2011; Nozue *et al*., 2013), while AtWRKY40 and AtWRKY63 repress or activate genes involved in high light signaling, respectively (Van Aken *et al*., 2013). Additionally, WRKY TFs have been implicated in nutrient deficiency response signaling pathways, where AtWRKY75 is induced by P_i_ starvation (Devaiah *et al*., 2007) and AtWRKY45 and AtWRKY65 are induced by carbon starvation (Contento *et al*., 2004). Likewise, previous studies in Arabidopsis (Col-0) revealed that WRKY1 expression is repressed by organic-N treatment (Gutierrez *et al*., 2008) and induced by N starvation (Krapp *et al*., 2011). Here, we describe i) how WRKY1 participates in genome-wide transcriptional reprogramming of Arabidopsis leaves in response to individual and combined light and nitrogen treatments, and ii) its potential role as an integrator of L and N signaling pathways toward the fine-tuning of plant energy status.

## RESULTS

### Gene regulatory network analysis reveals that WRKY1 is a hub in the nitrogen assimilation pathway

Our previous studies of N-regulatory networks in Arabidopsis identified a subnetwork of 367 connected nodes, including WRKY1 (Supplemental Data Set 1; Gutierrez et al 2008). In that initial network, protein:DNA interactions were predicted based on an overrepresentation of the regulatory motif for that transcription factor, and the expression of the transcription factor and putative target gene was highly (≥0.7 or ≤ −0.7) and significantly (*P* ≤ 0.01) correlated (Gutierrez *et al*., 2008). Subsequent Chromatin-IP analysis of a top hub (CCA1), in combination with bioinformatic CRE-motif analysis, revealed that the presence of a single binding site was sufficient for direct regulation of the target gene by the TF (Gutierrez *et al*., 2008). Based on these experimental results, we reanalyzed the N-response data in Gutierrez *et al*. (2008) by relaxing the predicted protein:DNA interaction to require a minimum of single regulatory motif for the TF in the promoter of a putative target, rather than an overrepresentation of binding sites. This resulted in an updated N-regulatory subnetwork, which increased the number of regulatory edges from WRKY1 to putative target genes (Supplemental Data Set 1). In this refined regulatory network, WRKY1 is one of the most highly connected TFs directly associated with metabolic genes involved in N assimilation, such as GDH1 (glutamate dehydrogenase), NIA1 and NIA2 (nitrate reductase 1 and 2), and ASN1 (asparagine synthetase), in which WRKY1 is predicted to activate GDH1, NIA1 and NIA2, and repress ASN1 (Supplemental Figure 1; Supplemental Table 1).

### WRKY1 target genes are involved in Nitrogen and Light signaling pathways

WRKY1 is predicted to be a major hub of an organic-N regulatory network (Gutierrez 2008), and is predicted to transcriptionally repress expression of ASN1, a gene regulated in response to light, nitrogen and carbon signaling (Thum *et al*., 2003). Here, we attempt to validate the involvement of WRKY1 as a hub of a nitrogen and light signaling network, by exposing *wrky1* mutant plants to N and L treatments. To test this, we compared three T-DNA alleles of WRKY1: *wrky1-1* (SALK_070989), *wrky1-2* (SALK_016954) and *wrky1-3* (SALK_136009) to wild-type Col-0 (the genetic background of the mutants). The *wrky1* mutant phenotype (SALK_016954) was previously described by (Qiao et al., 2016) in studies of its role in ABA signaling in stomata. In our studies of the three *wrky1* T-DNA mutant alleles, WRKY1 gene expression was altered from below the level of detection in *wrky1-1*, to 2% and 24% of WT WRKY1 expression levels in *wrky 1-2* and *wrky 1-3*, respectively (Supplemental Figures 2, 9-11).

To determine the effect of the *wrky1* T-DNA mutations on gene expression, *wrky1* plants were grown under “steady state” conditions, or in response to transient light and/or nitrogen treatments. For the “steady state” experiments, the three *wrky1* T-DNA alleles and Col-0 were grown on basal MS media under 16h/8h light/dark regime for 14 days. Shoot tissue was extracted for mRNA analysis by RT-qPCR and microarray analysis was performed with Affymetrix ATH1 array to identify genes mis-regulated in the *wrky1* mutants using Rank Product (Breitling *et al*. 2004) statistical analysis (See Methods). The “core set” of WRKY1 regulated genes were defined as those that are mis-regulated in the most severe knock-down mutant, *wrky1-1*, and *either* in *wrky 1-2* or *wrky 1-3*. This analysis identified 256 genes that are up-regulated and 117 genes that are down-regulated in the *wrky1* mutants (Supplemental Table 2) (Figure 1A). The 117 genes down-regulated in the *wrky1* mutants (i.e. genes induced by WRKY1) include a significant over-representation (pval <0.01) of genes enriched in GO-terms (BioMaps function in VirtualPlant 1.3; Katari *et al*., 2010) involved in secondary metabolic processes such as defense response and response to stress (Figure 1A). This role of WRKY transcription factors in the stress response has been reported for several members of the WRKY family of TFs (Qiao et al., 2016; Agarwal *et al*., 2011; Rushton *et al*., 2010; Chen *et al*., 2012). By contrast, the 256 genes up-regulated in the *wrky1* mutants (i.e. genes repressed by WRKY1) were significantly enriched for GO-terms involved in primary metabolic processes (pval=0.0001), response to carbohydrate stimulus (pval=2.3e-05), regulation of nitrogen compound metabolic process (pval=4.9e-05), and response to light stimulus (pval=0.0003) (Figure 1A). This result reveals new regulatory roles for WRKY1 as a transcriptional repressor involved in N and C-metabolism and light signaling.

**Figure 1.**
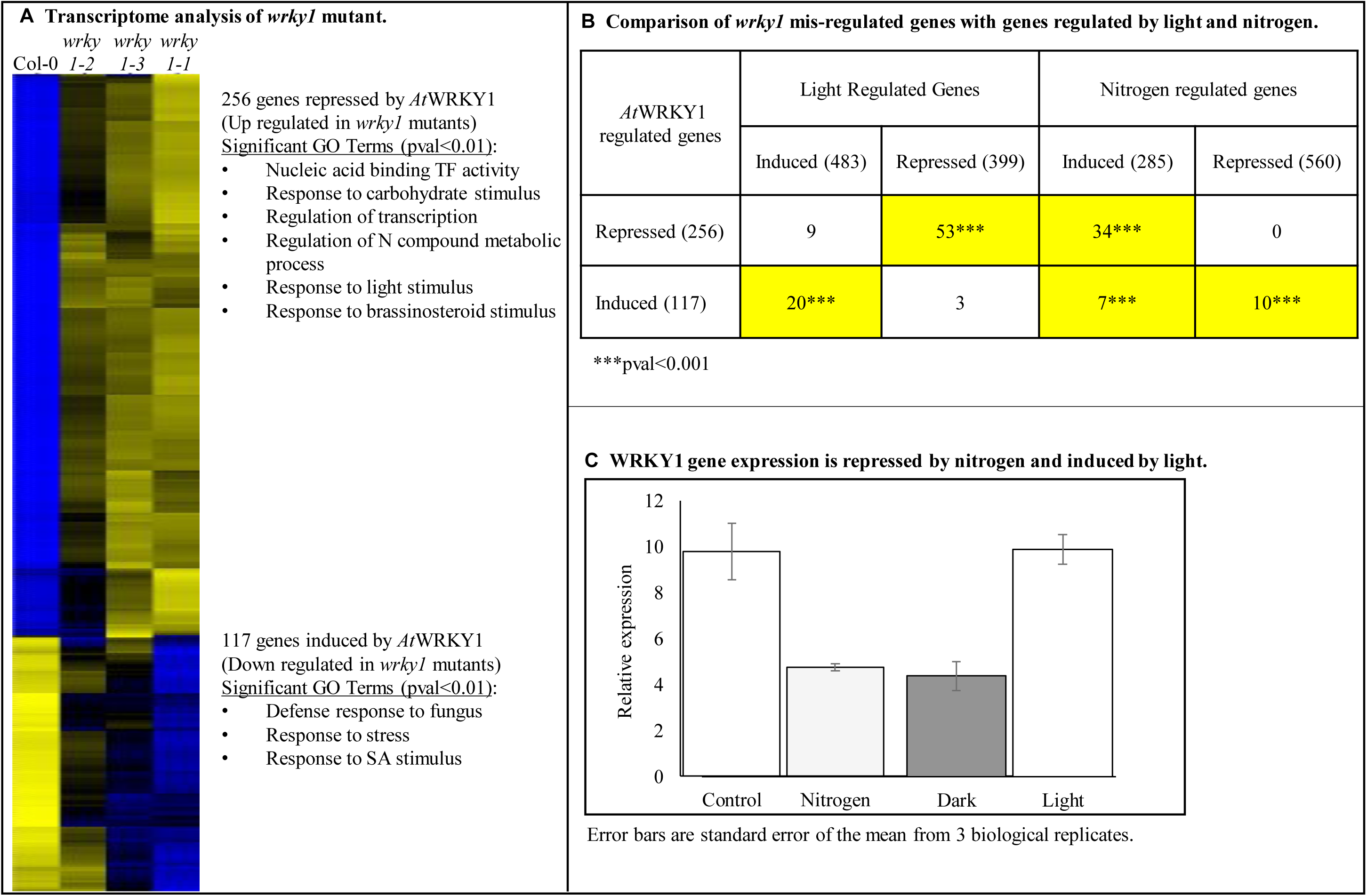
Down-regulation of WRKY1 results in mis-regulation of genes involved in light, nitrogen, and stress response pathways. **A.** Transcriptome analysis of WT (Col-0) and mutant *wrky1-1* (SALK_070989), *wrky1-2* (SALK_016954) and *wrky1-3* (SALK_136009) seedlings. Heat map of transcriptome data includes genes with significant (pval<0.01; FDR 5%) change in expression from WT in *wrky1-1* and *wrky1-2* or *wrky1-3*. Top significantly (pval<0.01) overrepresented GO terms are listed. B. Significance of overlaps (pval<0.001) of WRKY1 regulated, light regulated (Nozue et al., 2013), and nitrogen regulated (Gutierrez et al., 2008) gene sets were calculated using the GeneSect (R)script using the microarray as background. Total number of genes are inside parentheses; number of overlapping genes are shown in boxes. Boxes in yellow have p-value <0.001, indicating the size of the intersection is higher than expected. **C.** Relative expression levels of WRKY1 in WT (Col-0) seedlings in response to nitrogen (control is 20 mM KCl, treatment is 20 mM NH_4_NO_3_ + 20 mM KNO_3_^−^) and light treatments (control is normal long day, treatment is extended dark). Error bars are standard error of the mean; 3 biological replicates.

To identify the regulatory elements in the WRKY1 target genes, we performed a search for known cis-regulatory elements (CREs) in the putative promoter regions (2kb 5’ upstream of TSS sequences) of the genes mis-regulated in the *wrky1* mutants. We found a statistical over-representation of the W-box promoter motif (e-value = 5.17e-05) among the 117 genes down-regulated in the *wrky1* mutants. Although the W-box motif was present in nearly all of the genes up-regulated in the *wrky1* mutants (on average 2.18 W-box elements per promoter), it was not statistically over-represented. Instead, the I-box (e-value = 1.15e-72), GATA (e-value = 1.23e-46), ABRE-like (e-value = 3.27e-28), and G-box (e-value = 2.06e-22) motifs were the most statistically over-represented among the 256 up-regulated genes. The de-repression of these genes in the *wrky1* mutants could be due to an increase in factors that bind these other motifs. It is possible that these genes are either indirect targets of WRKY1 or that WRKY1 is part of a TF-complex that represses their expression in the wild type. Interestingly, a search for all known protein-protein interactions involving WRKY1, using the Arabidopsis Interactions Viewer (Geisler-Lee et al., 2007), revealed a single interaction (PSICQUIC confirmed by affinity chromatography) with General Regulatory Factor 1 (GRF1, AT4G09000), which is a G-box factor whose native form is as a hetero-dimer. So one hypothesis is that in the WT, WRKY1 interacts with GRF1 as a heterodimer to down-regulate the expression of these genes.

Investigation of the core genes involved in the N assimilation pathway revealed that down-regulation of WRKY1 expression affected the expression of genes involved in both nitrogen uptake and organic-N metabolism and catabolism. In WT plants, the expression of genes encoding several nitrate transporters as well as genes directly involved in glutamine biosynthesis are up-regulated, while genes involved in glutamine catabolism, such as ASN1 (asparagine synthetase1/DARK INDUCIBLE 6), are down-regulated during the light period. However, in *wrky1* mutant plants, the nitrate transporters NRT1.7 and NRT3.1 and the glutamate receptor GLR1.1 are down-regulated while ASN1 is up-regulated in the light. Thus, these transcriptome results provide support for the predicted edge between WRKY1 and ASN1 from the network analysis described above (Supplemental Figure 1). The observed overall reprogramming of the nitrogen network also indicates that WRKY1 is likely a regulatory molecule for this pathway.

The involvement of WRKY1 in mediating temporal responses to Light (L) and Nitrogen (N) signals was further investigated by intersecting the list of *wrky1* mis-regulated genes with lists of genes previously identified as responsive to N-treatments (Gutierrez *et al*., 2008) or L-treatments (Nozue *et al*., 2013) (Figure 1B). Genes normally repressed by WRKY1 in WT (i.e. the 256 genes up-regulated in the *wrky1* mutants) share a significant overlap with genes repressed by L-treatments (pval <0.001) (Figure 1B). By contrast, genes induced by WRKY1 in WT (i.e. the 117 genes down-regulated in the *wrky1* mutant) share a significant overlap with genes induced by L-treatments (pval <0.001) (Figure 1B). With regard to a potential role for WRKY1 in N-signaling, genes normally repressed by WRKY1 in WT (i.e. induced in the *wrky1* mutant) are induced by N-treatments (pval <0.001), while genes induced by WRKY1 (i.e. repressed in the *wrky1* mutants) are either repressed or induced by N-treatments (pval <0.001) (Figure 1B).

These reciprocal patterns of expression of genes regulated by WRKY1 in L and N treatment datasets introduce the hypothesis that WRKY1 is an integrator of L and N signaling pathways. This hypothesis is supported by the finding that WRKY1 expression is independently and reciprocally regulated by L and N treatments. Specifically, WRKY1 expression is induced by L-treatment, and repressed by N-treatment (Figure 1C). To further investigate the regulatory role of WRKY1 in N or L signaling, and the possible crosstalk, we exposed WT and *wrky1* mutants to three treatments: i) L treatment; ii) N treatment; and iii) combined N and L treatments, as described below (Table I). Three separate treatments were performed, as opposed to a single combined treatment, to eliminate omitted-variable bias and accurately determine the role of WRKY1 in the independent L and N signaling pathways and in regulating crosstalk between pathways, since transcriptional changes in response to double abiotic stress treatments are not predictable from responses to single stress treatments (Rasmussen *et al*., 2013; Prasch and Sonnewald 2013).

**Table 1.** Experimental design for light only (L), nitrogen only (N), and combined light and nitrogen (LN) treatments.

| Experiment | Light | Nitrogen |
| --- | --- | --- |
| L1 | + | + |
| L2 | - | + |
| N1 | + | - |
| N2 | + | + |
| LN1 | - | - |
| LN2 | - | + |
| LN3 | + | - |
| LN4 | + | + |

### WRKY1 mediates the light-repression of genes involved in organic resource catabolism

The finding that WRKY1 regulates genes implicated in light and nitrogen signaling (e.g. ASN1), inspired us to further examine the role of WRKY1 in the regulation of genes involved in the L response. This was investigated by comparing light-regulated gene expression in the null *wrky1-1* mutant, compared to WT Col-0 (Figure 2). For this experiment, seedlings were grown on MS media for 13 DAP and either maintained in normal light/dark cycle, or moved to extended darkness for 24 h prior to harvest. Two-way ANOVA of transcriptome data followed by FDR correction (pval<0.01 for the ANOVA model) identified 1,110 genes with a significant Light × Genotype interaction term (pval<0.01 for the coefficient of the term). Intersection of the 1,110 genes regulated by a LxG interaction with the 373 genes mis-regulated by knock-down of WRKY1 under “steady state” L-conditions (Figure 1), revealed a 35% overlap at a high-level of significance (pval<0.001). The large number of affected genes and highly significant overlap with genes mis-regulated in the *wrky1* mutant at steady-state conditions suggest a strong involvement of WRKY1 in L-signaling.

**Figure 2.**
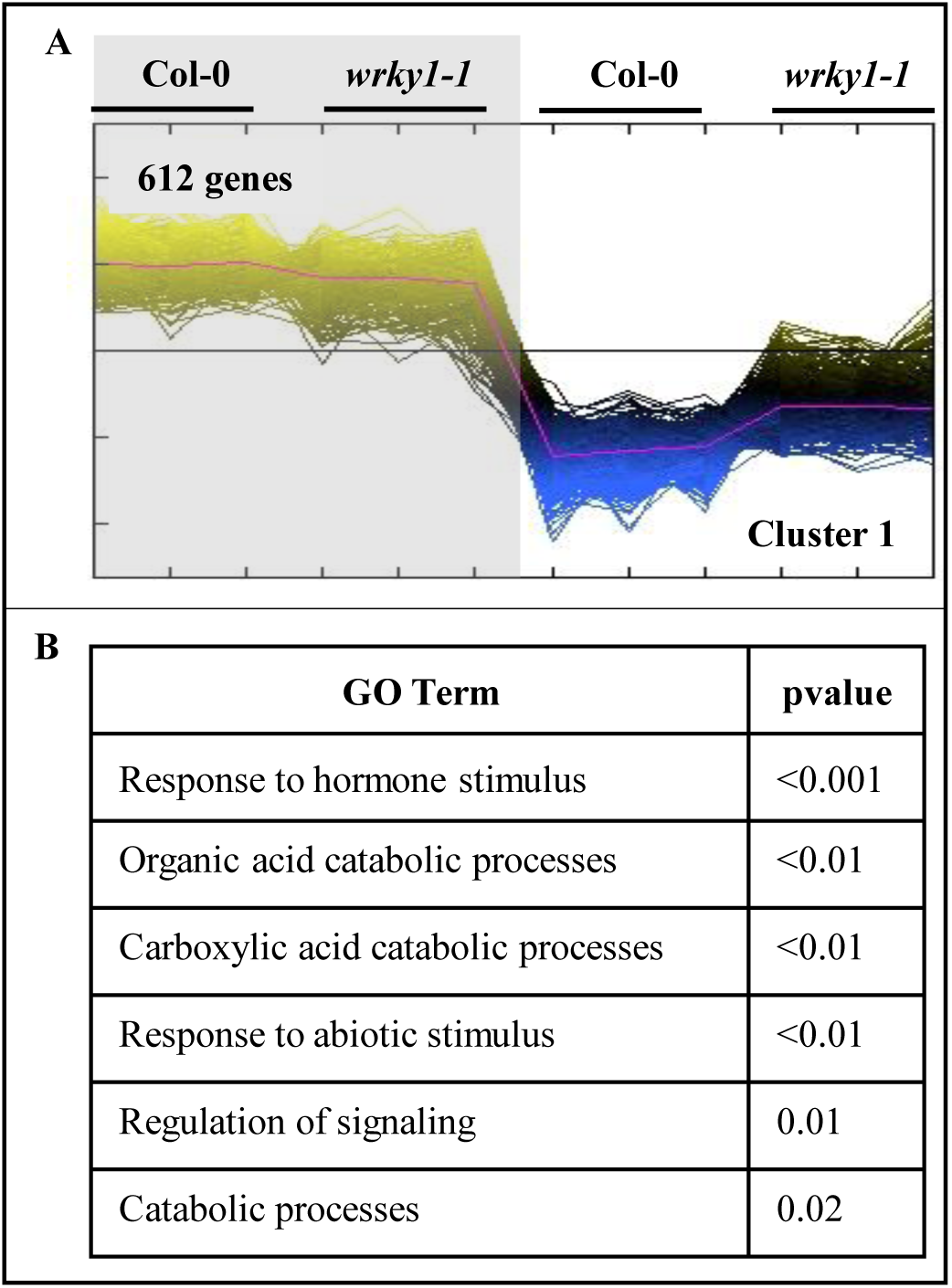
Cluster analysis of genes with significant Genotype × Light interaction reveal loss of light repression for some dark inducible genes. **A.** Cluster analysis of genes with significant (pval<0.02, FDR 5%) Genotype × Light interaction effect (1567 genes). Shaded area indicated dark conditions. Only Cluster 1 is shown, full cluster analysis can be viewed in Supplemental Figure 3. **B.** GO term analysis of gene cluster 1 with significant GxL effect.

To identify patterns of genes mis-regulated by light in the *wrky1* mutant, we performed cluster analysis of the microarray data. Specifically, Gene Expression Cluster analysis (Multiple Expression Viewer (MEV), QTC cluster analysis) and TukeyHSD analysis were performed on the genes with significant Light × Genotype interaction. Cluster analysis of the genes mis-regulated in response to a GxL interaction resulted in 11 distinct gene expression clusters (Figure 2; Supplemental Figure 3A). Cluster 1 is the largest (612 genes), which is a set of genes that have partially lost light-repression in the *wrky1* mutant (Figure 2A). For this set of genes, expression is normally repressed by L in the WT, but they are up-regulated in the *wrky1-1* mutant (Figure 2A). Genes in Cluster 1 include the dark-inducible genes DIN1, DIN4, DIN6/ASN1, and DIN10 (Fujiki *et al*., 2001). The intersection of Cluster 1 genes with previously identified light-induced or light-repressed genes (Nozue *et al*., 2013) revealed a significant overlap with the light-repressed genes (pval<0.001) (50 genes). Cluster 1 genes comprise GO-term enrichments (BioMaps) for organic acid and carboxylic acid catabolic processes (pval<0.01) (Figure 2B). This observation suggests that WRKY1 plays a large regulatory role in the light-repression of genes involved in catabolism of organic resources, which are specifically required in plants exposed to extended darkness. Two other highly significantly overrepresented GO-terms are for “response to abscisic acid stimulus” (p-value = 0.01) and “regulation of abscisic acid mediated signaling pathway” (p-value = 0.07). These responses are consistent with recent studies that showed a role for WRKY1 in ABA signaling in response to drought (Qiao et al., 2016).

Unique and significant GO-term enrichments (BioMaps) were also uncovered for the other clusters of genes regulated by a Light × Genotype interaction in the *wrky1* mutants (Figure 2; Supplemental Figure 3B). These include N-compound metabolic processes (Cluster 2), disaccharide biosynthetic processes (Cluster 4), generation of precursor metabolites and energy (Cluster 5), ATP biosynthetic process (Cluster 10), and carbohydrate metabolic process (Cluster 11).

Finally, TukeyHSD analysis of the Light × Genotype interaction term, revealed a larger Genotype effect in the Light (901 genes) than in the Dark (499 genes) (Tukey pval<0.01), while the Light effect was 89% similar between WT and the mutant. These results suggest a significant role for WRKY1 in the regulation of light responsive genes (Figure 2, Supplemental Figure 12).

To summarize, in the light, WRKY1 specifically: i) *represses* a network of genes that are required to catabolize cellular resources when light (i.e. in the form of carbon) is limited (Cluster 1), and ii) *activates* a subset of genes involved in the biosynthesis of energy-dependent metabolites synthesized during the day (Clusters 4 and 5) (see summary Figure 7A). By contrast, in the dark, WRKY1 i) *activates* a subset of genes involved in processes of respiration and the production of energy metabolites, and ii) *represses* genes involved in energy expensive, secondary metabolic processes (see summary Figure 7A).

### WRKY1 mediates transcriptional reprogramming in response to N treatment

The steady-state analysis of the *wrky1* mutant revealed mis-regulation of genes involved in the N-assimilation pathway. However, to test if WRKY1 is involved in the regulation of plant responses to N-signaling, it was necessary to explore changes in expression of genes involved in the N signaling pathway to a transient N treatment in both the most severe *wrky1-1* T-DNA mutant (SALK_070989) and WT (Col-0) seedlings. To do this, *wrky1-1* and WT seedlings were grown on basal MS media supplemented with 1mM KNO_3_^−^ for 14 DAP under long day cycle. At the start of day, seedlings were transferred to either 60mM N (20mM KNO_3_^−^ + 20mM NH_4_NO_3_) or 20mM KCl (control) for two hours prior to harvest. A two-way ANOVA of genome-wide transcriptome data followed by FDR correction of the ANOVA model (pvalue<0.01) uncovered 123 genes with a significant Nitrogen × Genotype interaction term (pvalue<0.02). Of these 123 genes, 11 had a significant overlap (pval<0.05) with N-regulated genes in a N-regulatory network previously identified by Gutierrez *et al*. (2008), including nitrate reductase 1 (NIA1). This result indicates that a different network of genes responds to transient N treatment when WRKY1 is absent than when it is present.

To better understand the biological role of the 123 genes whose expression is regulated by WRKY1 and show a significant NxG interaction, Gene Expression Cluster analysis (QTC function in MEV) and TukeyHSD analysis was performed. First, cluster analysis resulted in six distinct gene expression clusters (Figure 3A), involved in different biological processes (Figure 3B). Clusters 1 and 3 contained genes with the most significant over-representation of GO terms (BioMaps), in which genes in Cluster 1 are predominantly involved in Cellular Homeostasis (pval=0.0009), while genes in Cluster 3 are involved in Translation (pval=2.7E-11) and Cellular protein metabolic process (pval 7.89E-07) (Figure 3B). Further inspection of the gene expression clusters 2 and 3 revealed that a subset of genes do not respond to N-limitation in the mutant like they do in plants with WT WRKY1 function. This finding suggests that the role of WRKY1 in the N-signaling pathway may be as a transcriptional regulator in the control of the N starvation signaling processes. According to this hypothesis, the subset of genes must meet two criteria: i) in the *presence* of N, gene expression is the same between the WT and *wrky1-1* mutant; and ii) in the *absence* of N, gene expression is different between the WT and *wrky1-1* mutant. To identify a full subset of genes that meet these criteria, we compared TukeyHSD results from the two-way interaction terms for “N effect in *wrky1-1* mutant plants” and for “Genotype effect in the absence of N”. This analysis uncovered 38 genes that met our criteria, of which 18% are involved in carbon compound and carbohydrate metabolism (including O-Glycosyl hydrolases family 17 protein (At5g58090); UDP-Glycosyltransferase superfamily protein (At2g36970); Phosphofructokinase family protein (At1g20950), and XTH6_xyloglucan endotransglucosylase/hydrolase 6 (At5g65730) and others (see Supplemental Table 4). This result provides further support that WRKY1 is involved in N and energy related signaling pathways. Additionally, TukeyHSD analysis revealed that the N response was significantly altered in the *wrky1* mutant for 63 genes (pval<0.01). Of these, 43 genes responded to N in WT but not in the *wrky1-1* mutant, while 18 genes had a significant N response only in the *wrky1* mutant seedlings.

**Figure 3.**
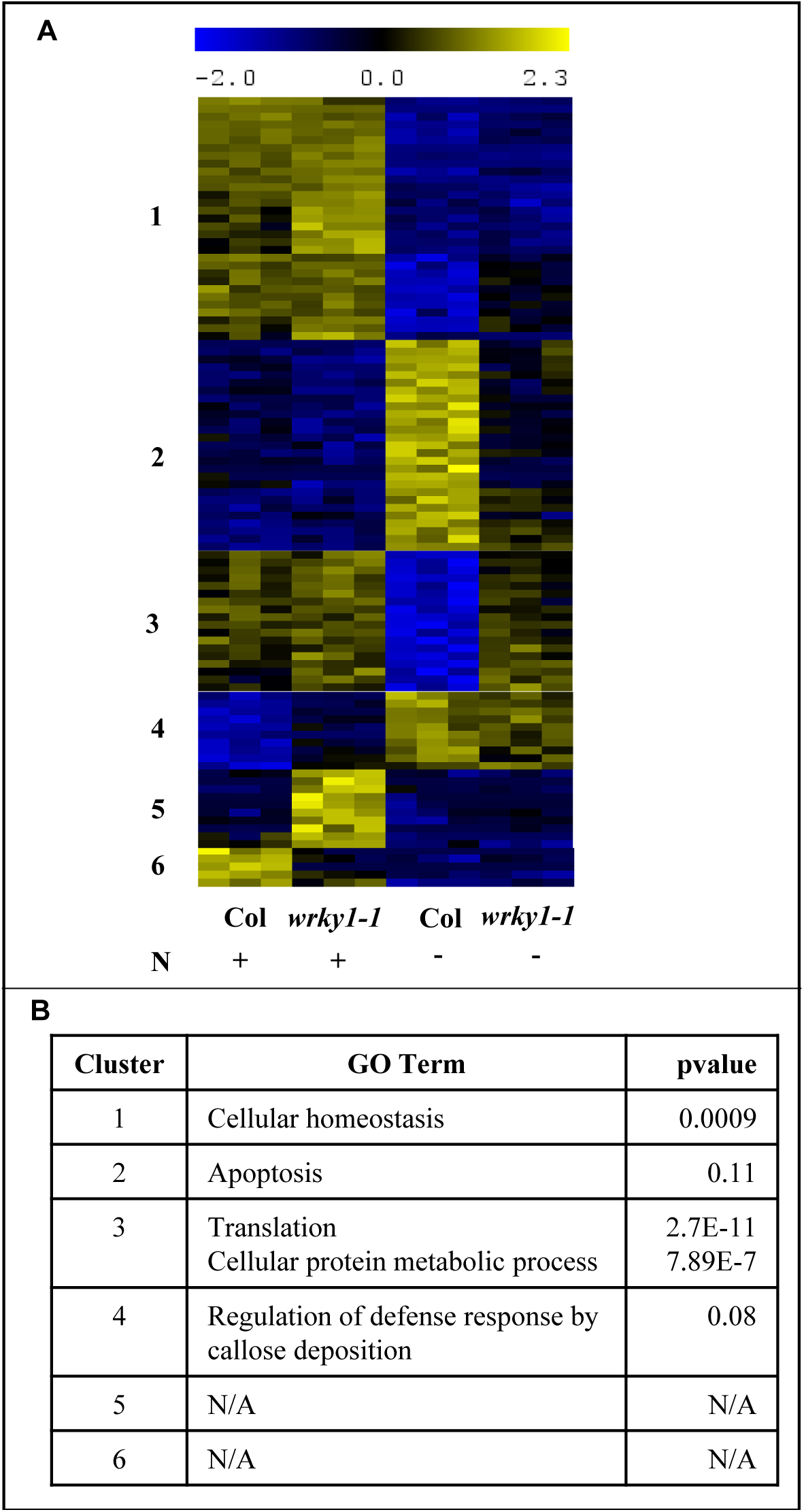
Cluster analysis of genes with significant Nitrogen × Genotype effect reveal that WRKY1 participates in plant response to N-limitation. **A.** Cluster analysis of genes with significant (pval<0.02, FDR 5%) Genotype × Nitrogen interaction effect (123 genes). **B.** GO term analysis of gene clusters with significant GxN effect.

Cumulatively, these analyses reveal that: i) WRKY1 regulates a different transcriptional program of genes in response to transient N treatment compared to steady-state N conditions. ii) Fewer genes are mis-regulated by knock-down of WRKY1 gene expression in response to transient N treatment compared to steady-state N conditions. iii) N response is altered in *wrky1* mutants compared to WT, in which WRKY1 represses genes involved in defense response in the presence of N. However, in the absence of N, WRKY1 activates genes involved in apoptosis (Cluster 2, Figure 3) and represses genes involved in translation and protein metabolic processes (Cluster 3, Figure 3) that require N (Figure 7B). This last result further supports the hypothesis that WRKY1 is involved in energy conservation, where under N limiting conditions WRKY1 activates genes involved in recycling of cellular resources while simultaneously suppressing genes involved in energy expensive protein biosynthesis.

### Combined Light and Nitrogen treatments reveal that WRKY1 regulates crosstalk between L and N signaling pathways

The above studies collectively support the hypothesis that WRKY1 is a regulatory node in the Light and Nitrogen response network in Arabidopsis. This implication along with evidence from previous research (Jonassen *et al*., 2008; Nunes-Nesi *et al*., 2010; Obertello *et al*., 2010; Krouk *et al*., 2009) reinforce the notion that transcriptional crosstalk occurs between light and nitrogen signaling pathways to fine-tune plant energy status. This hypothesis was further investigated by performing combined treatments with L and N on WT (Col-0) and *wrky1-1* null mutant (SALK_070989) seedlings. Here, the aim was to determine if combined L and N treatments will reveal different transcriptional reprogramming by WRKY1 than is observed in response to individual L or N treatments, as has been observed for ten Arabidopsis ecotypes in response to single and double stress treatments (Rasmussen *et al*., 2013).

To test this hypothesis, seedlings were grown on basal MS media supplemented with 1 mM KNO_3_^−^ under long day light cycle for 13 days. For dark treatment, WT and mutant seedlings were moved to continuous dark for 24 h prior to N treatment. For N treatment, seedlings were transferred to basal MS media supplemented with 60 mM N (20 mM KNO_3_^−^ plus 20 mM NH_4_NO_3_) at start of light cycle (or putative light cycle for dark treated seedlings) for two hours. Shoot tissue was extracted for expression analysis by microarrays, while data was analyzed by three-way ANOVA followed by FDR correction of the ANOVA model (pval<0.01). Three-way ANOVA revealed significant main effects, two-way interaction effects, and a three-way interaction effect (Table 2), which identified 724 genes with significant three-way interaction of Genotype, Light and Nitrogen. Gene network analysis was used to organize these 724 WRKY1- dependent genes into a network, revealing predicted interactions among nodes based on co-expression and protein:DNA regulatory interactions (Supplemental Figure 4).

**Table 2.** Results of three-way ANOVA, for individual and interaction terms. Genotype (G), nitrogen (N), light (L). No. Genes is the number of ATH1-genechip identifiers (probes).

| Effect | No. Genes |
| --- | --- |
| **G** | **2356** |
| **N** | **5062** |
| **L** | **10158** |
| **GxN** | **1022** |
| **GxL** | **1459** |
| **NxL** | **2114** |
| **GxNxL** | **700** |

To identify groups with similar expression patterns within the 724 genes whose expression is affected by a Genotype × Light × Nitrogen interaction, cluster analysis was performed (QTC function in MEV) for genes with a significant three-way interaction term (pval<0.01), which resulted in eight distinct clusters (Figure 4A). GO term analysis (BioMaps) revealed unique and significant biological functions for genes in clusters 1-6 and 8 (Figure 4B), including response to light stimulus (Cluster 1, pval=0.04); photosynthesis (Cluster 2, pval 2.7E-11); embryo development (Cluster 4, pval 3.8E-8); response to nitrate (Cluster 5, pval 0.0003); and regulation of hormone levels (Cluster 6, pval=0.02). Gene network analysis resulted in a network in which genes were grouped by significant (pval<0.05) over-representation of shared biological processes (BinGO Plug-in Cytoscape) (Figure 5). The largest gene clusters had over-represented GO terms for “metabolic processes” (177 genes); “response to stimulus” (106 genes); and “developmental process” (60 genes) (Figure 5). This analysis provides insight into the biological processes influenced by the crosstalk between Nitrogen and Light signaling pathways in which WRKY1 is a regulatory node.

**Figure 4.**
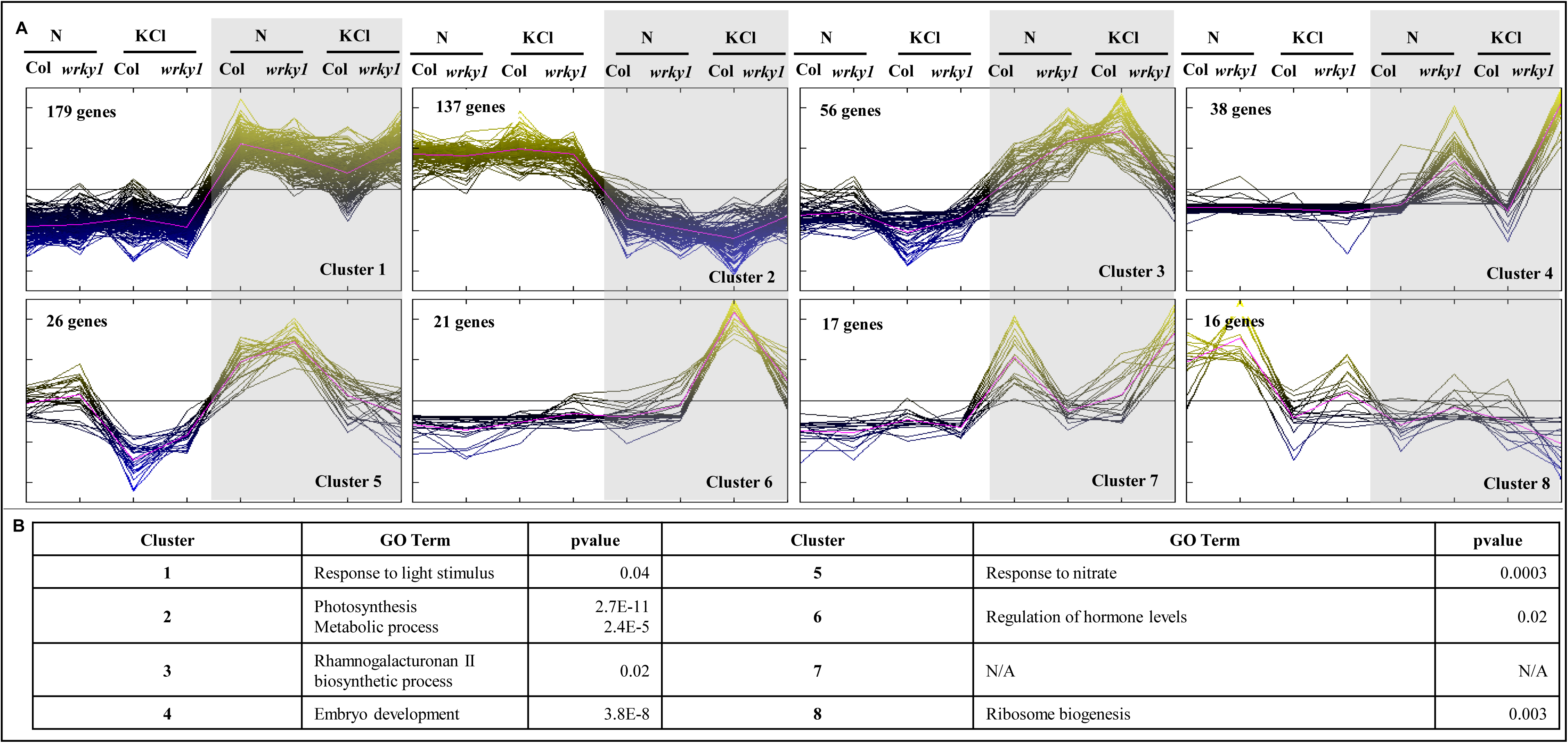
Combinatorial treatment of *wrky1* mutants and WT with Nitrogen and Light results in a significant 3-way GxLxN interaction. Cluster analysis of genes with significant (pval<0.01, FDR 5%) Genotype × Nitrogen × Light interaction effect (724 genes). **B.** GO term analysis of gene clusters with significant GxNxL effect. Shaded area indicated dark conditions. N = Nitrogen treatment; KCl = control treatment; Col = Col-0; *wrky1 = wrky1-1*.

**Figure 5.**
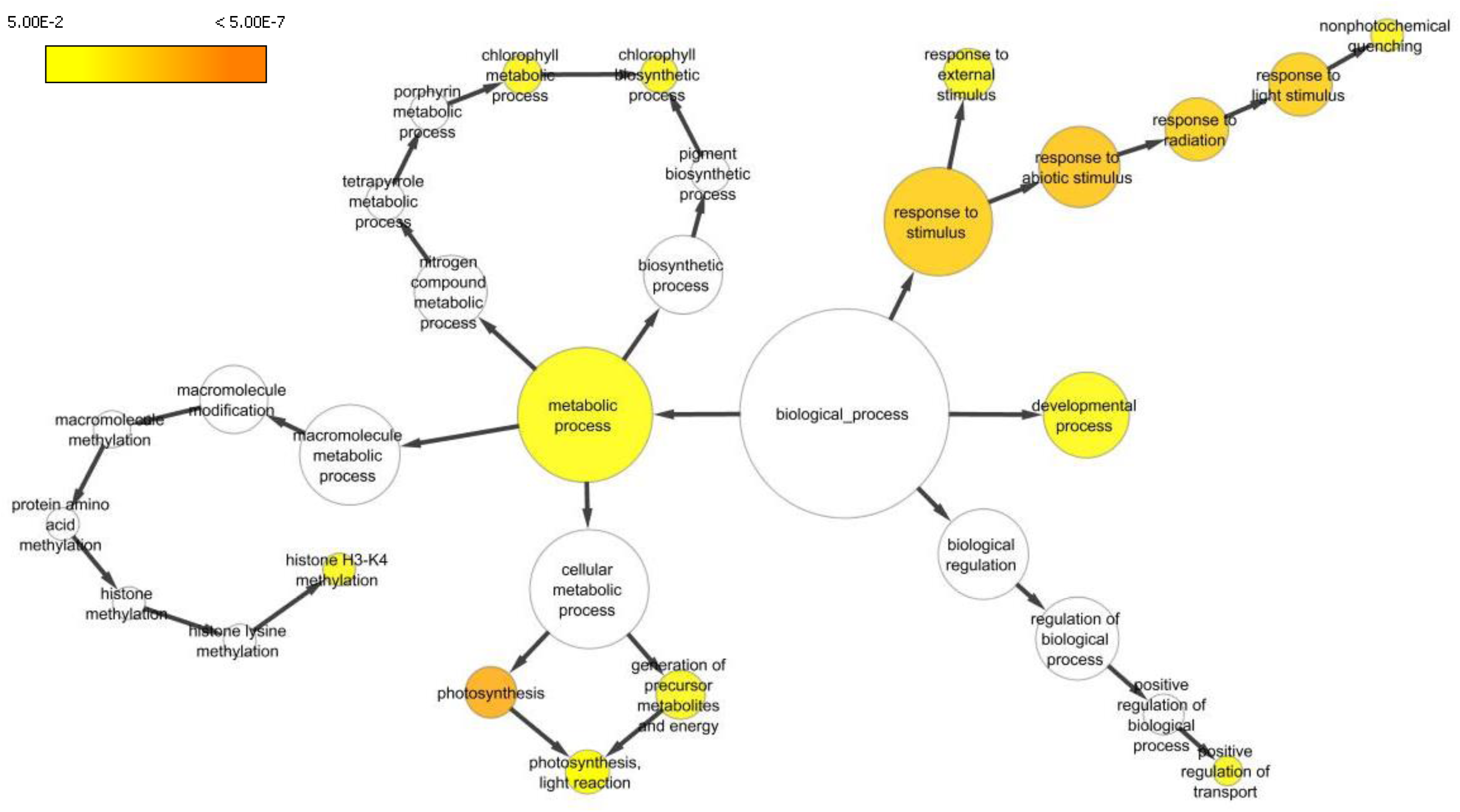
Genes commonly regulated by WRKY1, Nitrogen, and Light comprise a network of diverse biological functions. Network of statistically overrepresented GO terms in the set of 724 genes with significant G × N × L interaction effect. Node area is proportional to the number of genes within the functional category (i.e. more genes equals larger node). Colored nodes are significantly overrepresented (see color legend), while white nodes are not significantly overrepresented (BinGO, Maere et al., 2005).

To fully interpret the three-way interaction term (LxNxGenotype), genes with significant three-way interaction were further investigated to statistically determine how the various two-way interactions differed across the levels of the third variable using a sequential ANOVA approach (Figure 6). Principle component analysis (PCA) of all single and combined treatments revealed that light was the dominant effect (PC1), accounting for 49% of the variance, while N corresponded to PC2, explaining 30% of the total variance (Supplemental Figure 5). Therefore, the first iteration of sequential ANOVA was performed across levels of the Light variable, and the second iteration across levels of the Nitrogen variable. Two-way ANOVAs of genes with significant three-way interaction term under each light condition revealed significant NxG interaction exclusively in the DARK for 78% of genes (pval<0.05); exclusively in the LIGHT for 12% of genes; and in both DARK and LIGHT conditions for 10% of genes (Figure 6).

**Figure 6.**
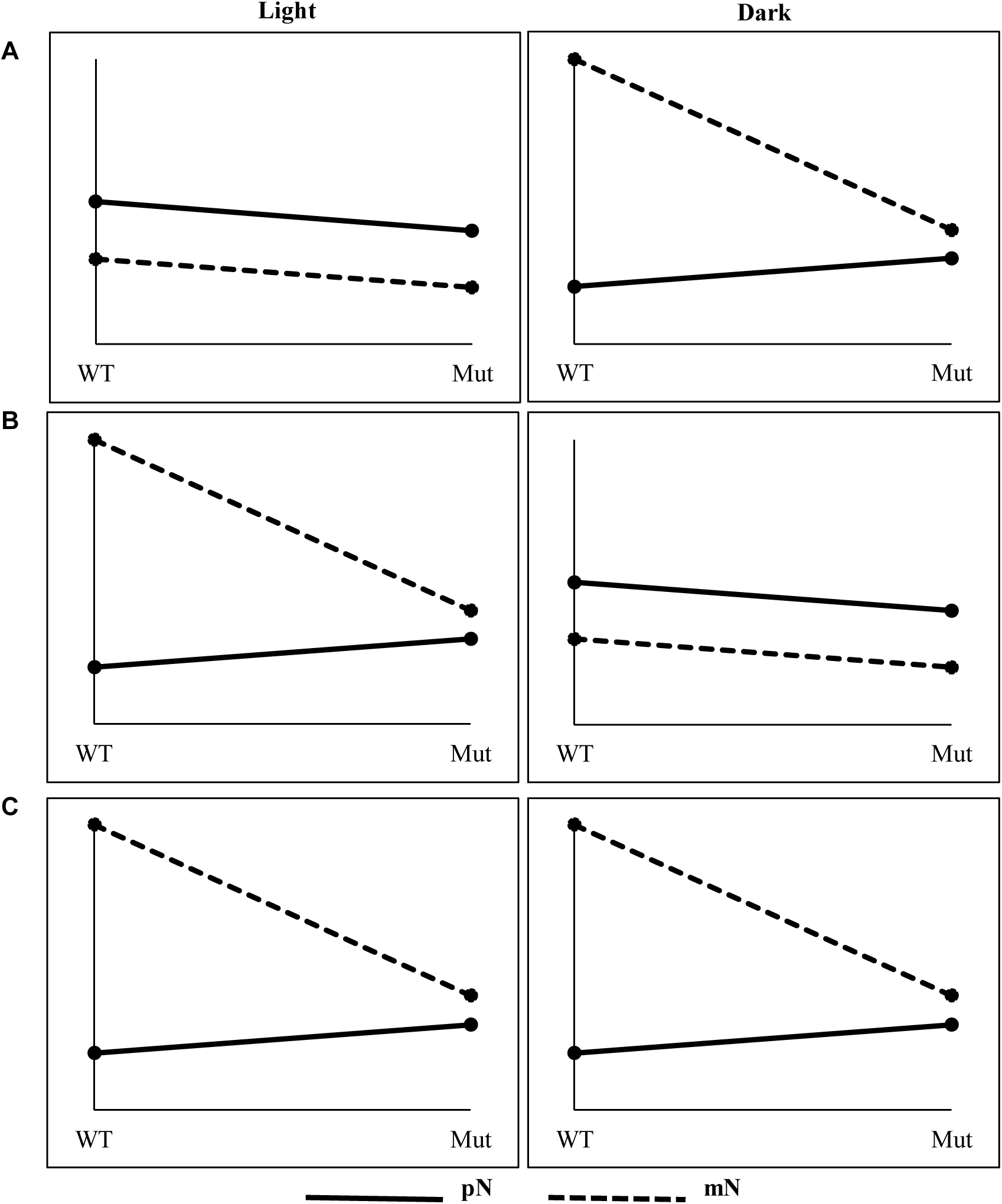
Graphical representation of the sequential ANOVA to interpret the 3-way interaction term. These plots represent the general observations for the 724 genes with significant three-way interaction. This example shows the three scenarios we observed for how gene expression changes in the light and the dark in response to the nitrogen variable across levels of the genotype variable. In our study, 78% of the genes only have a significant G × N interaction in the Dark (**A**), 12% of the genes only have a significant G × N interaction in the Light (**B**), and 10% of the genes have a significant G × N interaction in both Dark and Light conditions (**C**). WT = Col-0, Mut = *wrky1-1*.

These results indicate that the interaction between Genotype and N are most significant in DARK conditions, which was also observed visually from the cluster analysis (Figure 4). Genes with significant NxG interaction in the DARK were uniquely and significantly enriched in GO terms (BioMaps) for photosynthesis (pval=0.002), response to light stimulus (pval=0.004), and glutamine metabolic process (pval=0.05). Alternatively, genes with significant NxG interaction exclusively in the LIGHT (12% of genes with significant three-way interaction) were uniquely enriched in GO terms for lignin metabolic process (pval=0.02). Genes with significant two-way interaction term (pval<0.05) in both LIGHT and DARK conditions were enriched in GO terms for mRNA catabolic process (pval=0.04) and intracellular transport (pval=0.04).

To better understand the two-way NxG interaction term, the next step in the sequential ANOVA analysis was to perform one-way ANOVAs under each N condition while holding the light variable as either “DARK” or “LIGHT”. Here, we focus on one-way ANOVA results in DARK conditions since it was revealed as the dominant effect from the previous step. A comparison of one-way ANOVA models of genes with significant two-way interaction term in the dark revealed that there was a significant Genotype effect for 16% of genes exclusively in the *presence* of N, 47% of the genes exclusively in the *absence* of N, and 37% of genes in both presence and absence of N.

The genes with significant Genotype effect in the DARK exclusively in the *presence* of N have significant enrichment of GO terms for S-glycoside catabolic process (pval=0.04) and carbohydrate catabolic process (pval=0.05), where in response to transient N treatment, WRKY1 activates a network of genes involved in the remobilization of cellular carbon resources and represses genes involved in biogenesis (Figure 7C; Supplemental Figure 6). However, genes with significant Genotype effect in the DARK exclusively in the *absence* of N are uniquely enriched in GO terms for light stimulus (pval=0.0004); photosynthesis (pval=0.0016); and amine metabolic processes (pval=0.0088), in which WRKY1 activates genes involved in metabolic and biosynthetic processes for production of glutamine, tryptophan and chorismate, and represses genes that respond to light stimulus (Figure 7C). Additionally, genes with significant Genotype effect (pval<0.05) under both N regimes are enriched in GO terms for chlorophyll biosynthetic process (pval=0.004); reproductive process (pval=0.006); and embryo development ending in seed dormancy (pval=0.02).

**Figure 7.**
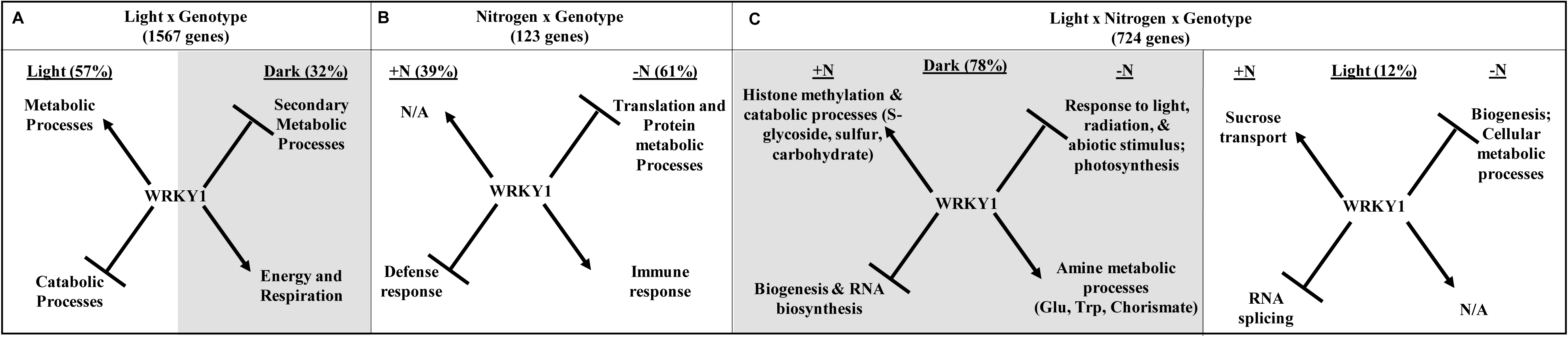
WRKY1 regulates genes in Light and Nitrogen pathways and is an integrator of Light and Nitrogen signaling. Putative mechanism by which WRKY1 regulates different transcriptional programs under three conditions: A. Light treatment; B. Nitrogen treatment; C. Combined Light and Nitrogen treatment. The most significantly overrepresented GO terms for biological process are shown. Arrows indicate activation, lines and bars indicate repression. Percentages are the number of genes from a given group that adhere to the proposed mechanism in each panel. Shaded areas indicate dark conditions. +N = nitrogen treatment; −N = control treatment.

These results indicate that the Genotype effect is weakest in the presence of N. This finding supports our earlier hypothesis that a less significant Genotype effect is observed between WT and *wrky1* mutants when N is present. Ultimately, the sequential ANOVA analysis can be interpreted that the Genotype effect caused by mutation of WRKY1 is revealed most significantly in the DARK and in the *absence* of N (Figure 7C; Supplemental Figure 6). This finding suggests a mechanism by which WRKY1 regulates a transcriptional program of genes in response to C and N limitation. Moreover, analysis of the three-way interaction term provides support that WRKY1 is a regulatory node connecting Nitrogen and Light signaling pathways. In this model of transcriptional regulation, WRKY1 modulates the expression of a new network of genes in response to simultaneous N and L signals compared to the transcriptional programs controlled by WRKY1 in response to either N or L signaling alone (Figure 7 A-D).

### Phenotypic analysis reveals the regulatory role of WRKY1 in nitrogen metabolism

Our transcriptional and bioinformatics analysis suggests a role for WRKY1 in the regulation of L and N signaling. To investigate the physiological effect of the *wrky1* mutation on plants, *wrky1-1* mutant and WT lines were grown on soil under low (0 mM N supplement) and high (50 mM N supplement) nitrogen fertilization regimes and subject to 14 days of growth on normal long-day light cycle. Plants grown for 14 days were then harvested for elemental analysis to assess total C and N. Total N analysis revealed a N × genotype effect (Figure 8A) in which there was more total N in the WT compared to the *wrky1-1* mutant only under low N conditions (p-value = 0.114) compared to high N conditions (p-value = 0.841). Likewise, there was a N × genotype effect for total C content (Figure 8B) where there was higher C content in the WT compared to the *wkry1-1* mutant under low N conditions (p-value = 0.102), and there was similar C content between WT and *wrky1-1* mutant under high N content (p-value = 0.785).

**Figure 8.**
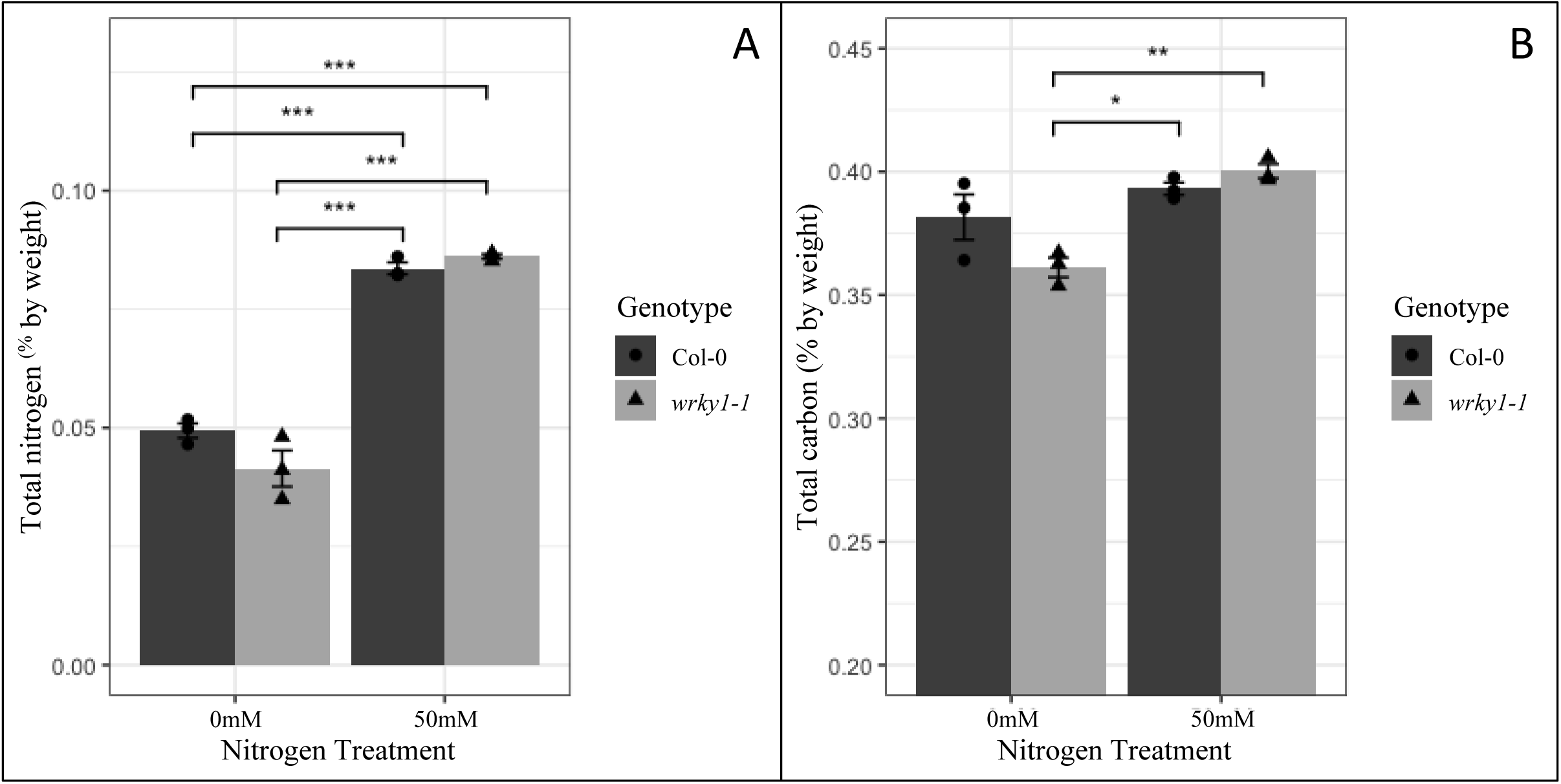
Total N and C contents in WT and *wrky1-1* plants. A. Total nitrogen by percent weight (mean ± SD % by weight) in Col-0 and wrky1-1 under 0mM and 50mM N supplement. B. Total carbon by percent weight (mean ± SD % by weight) in Col-0 and wrky1-1 under 0mM and 50mM N supplement. * and + indicates statistical significance as determined by two-tailed T-test: * = p-value < 0.05, ** = p-value < 0.01, *** = p-value < 0.001.

We further investigated the underlying metabolism by analyzing free amino acids and carbohydrates using GC-MS in an attempt to determine the underlying cause for the change in total N and C content in mutant compared to WT plants. Plants were grown for two weeks on MS media supplemented with either 0.5 mM KNO_3_^−^ or 10 mM KNO_3_^−^ then harvested. The majority of free amino acids were not significantly different between WT and *wrky1-1* mutant plants (Supplemental Figure 8). However, there was a higher concentration of glutamine in *wrky1-1* mutants under both low (p-value = 0.33) and high [NO_3_^−^] (p-value = 0.50) compared to the WT, and there was a slightly higher accumulation of aspartate in WT than mutant under low [NO_3_^−^] (p-value = 0.15) (Figure 9A-B). Upon examining carbohydrates, the *wrky1-1* mutant had higher concentrations of sucrose (Figure 9C) and its products glucose and fructose under both low (p-value = 0.36) and high (p-value = 0.16) NO_3_^−^ conditions. However, WT plants had higher concentrations of the dicarboxylic acid malate only under low NO_3_^−^ conditions (Figure 9D) (p-value = 0.06). These reciprocal patterns of glutamine/aspartate and sucrose/malate suggest a reprogramming of central C and N metabolism in *wrky1* mutant plants that result in lower overall C and N content when N is limiting. The WT function of WRKY1 may be to regulate genes involved in the redirection of flux through the TCA cycle away from glutamine biosynthesis and toward malate/asp synthesis under N-limiting conditions as part of a resource conservation mechanism.

**Figure 9.**
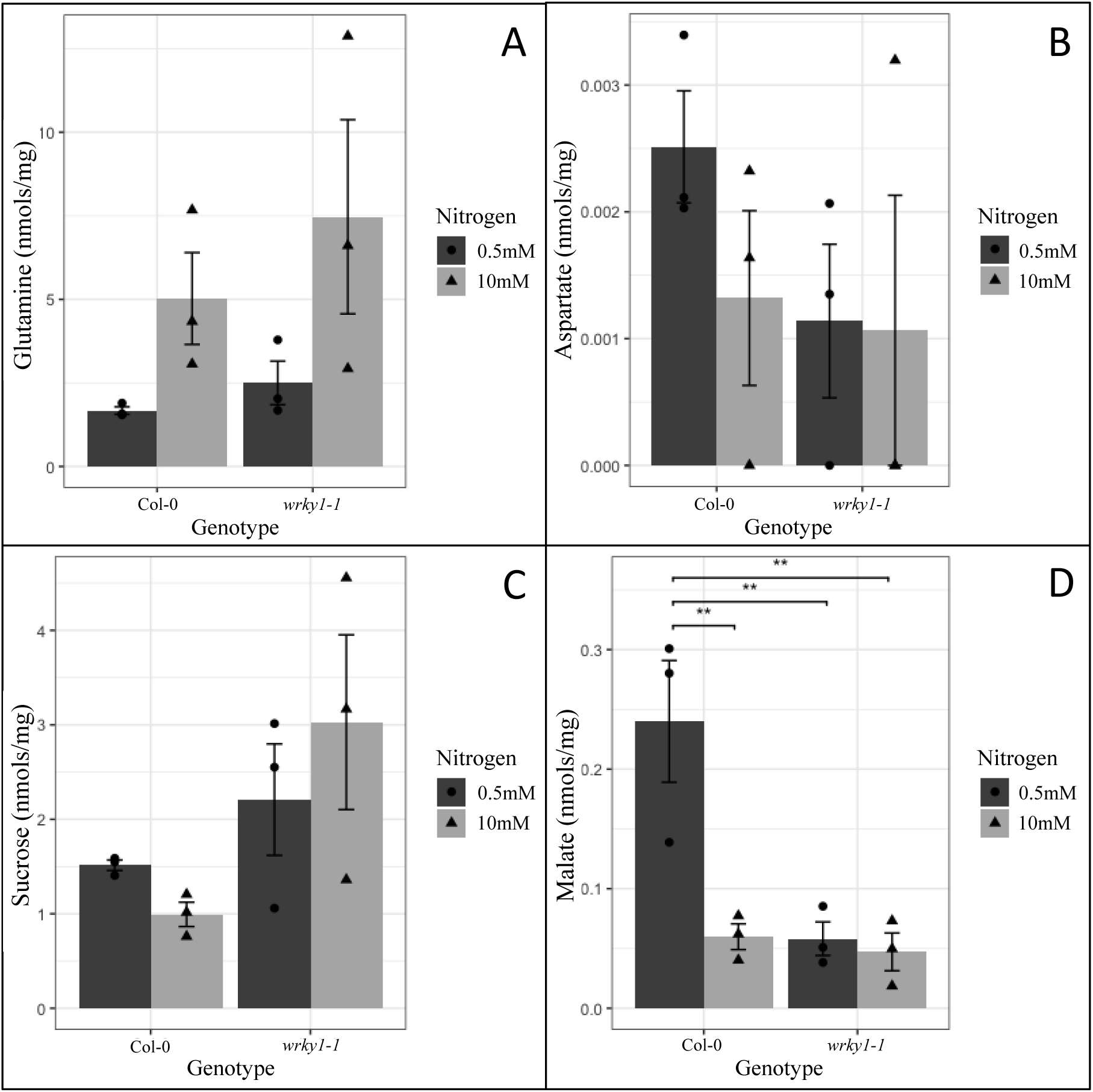
Measured metabolite levels (mean ± SD nmols/mg) in Col-0 and wrky1-1. A. glutamine; B. aspartate; C. sucrose; and D. malate, under nitrogen treatment of 0.5mM KNO_3_^−^ and 10.0mM KNO_3_^−^. * and + indicates statistical significance as determined by two-tailed T-test: + = p-value < 0.02, ++ = p-value < 0.01, * = p-value < 0.005.

## DISCUSSION

### WRKY1 regulates different transcriptional programs depending on the signal or combination of signals perceived

Our data on the *wrky1* T-DNA mutants reveal a defect in genome-wide expression that is dependent on light (L), nitrogen (N) and a combination of LxN. This suggests that WRKY1 mediates crosstalk between L and N signaling. Vert and Chory (2011) established two criteria for crosstalk to exist between two signaling pathways: i) “the combinatorial signal from both pathways should produce a different response than that triggered by each pathway alone”, and ii) “the two pathways must be connected directly or indirectly.” PCA analysis of the *wrky1* mutant revealed that light is the dominant effect among single and combined treatments (Supplemental Figure 4). However, to explain variance in gene expression, the ANOVA analyses revealed that a different transcriptional program is activated in response to concurrent N and L treatments in *wrky1* mutant and WT seedlings, compared to transcriptional response to individual N and L treatments. In addition, approximately 80% of the 724 WRKY1 regulated genes shared between the L and N pathways (those with significant three-way interaction term) are unique to the combined L and N treatment compared to individual treatments, indicating a direct connection between these pathways.

Our results for WRKY1 are similar to those reported by a recent study that revealed that 61% of transcriptional changes in ten Arabidopsis ecotypes in response to double abiotic stress treatments were not predictable from responses to single stress treatments (Rasmussen *et al*., 2013). Therefore, it is reasonable to suggest that WRKY1 is a regulator of crosstalk between L and N signaling pathways. Moreover, the resulting crosstalk network contains 29 transcription factors (Supplemental Table 3), of which 15 have significant (pval=5.37e-10) overrepresentation of the GO term “regulation of nitrogen compound metabolic process,” further indicating a direct connection between N and L signaling pathways. The sequential ANOVA performed here was able to effectively deconstruct the three-way interaction term to reveal dominant effects among the interaction to define the plant status under which WRKY1 mediates crosstalk between N and L signaling pathways.

Our analysis revealed a potential mechanism by which WRKY1 functions to repress genes involved in plant response to light stimulus and activate genes involved in amine metabolic processes when both light and exogenous N are limiting (Figure 7C). This example may be extrapolated as a mechanism by which plant transcription factors influence the partitioning of cellular resources in response to complex environmental signals. The intensive statistical analysis presented here can be used to decipher multifaceted interactions arising from similar or even more complex combinatorial experiments.

### WRKY1 is likely involved in an energy conservation mechanism in response to low energy signaling

ANOVA revealed that the majority of the 724 genes mis-regulated in the *wrky1-1 null* mutant had significant GxN interaction in the dark compared to the light. This result in combination with GO term analysis generates the hypothesis that WRKY1 is part of an energy conservation mechanism by which targets of WRKY1 remobilize C resources in the dark when N is abundant, but up-regulates N metabolism in the dark when N is limiting. In this mechanism, WRKY1 integrates information about cellular N and energy resources to trigger processes necessary for plant metabolism in response to a transient N-signal. For example, when both light (i.e. carbon) and nitrogen resources are limiting, genes involved in light response and photosynthesis were significantly up-regulated in the *wrky1* mutant. By contrast, genes involved in glutamine and tryptophan metabolic and biosynthetic processes were significantly down-regulated in the *wrky1* mutant under limiting conditions of light and nitrogen (Figure 7C). These results are supported by research by Urbanczyk-Wochniak and Fernie (2005), who uncovered the surprising result that several amino acid pools, including arginine, asparagine, and glutamate, are higher under N-deficient conditions compared to N-saturated conditions. Specifically, the authors discovered that N-deficient plants in low light conditions have increased carbohydrate content, and that glutamate and tryptophan metabolite pools increase initially in response to N-deficiency in both high and low light conditions. Our model of WRKY1 mediated regulation of genes in the dark provides transcriptional support for the observed changes in metabolite pools in response to N-deficiency in low light conditions (Urbanczyk-Wochniak and Fernie, 2005). In our own experiments, we observed that under normal light conditions but with N-limitation, there is a decrease in total N and C content in the *wrky1* mutant, but a higher concentration of glutamine and lower concentration of malate and aspartate compared to the WT (Figure 9 B and D). This analysis of free metabolite pools suggest that the *wrky1* mutant fails to redirect metabolism toward aspartate biosynthesis and instead maintains glutamine biosynthesis even when C and N resources are limiting. In future studies, it would be interesting to examine the amino acid content in proteins (protein-bound aas) to enhance our understanding of the underlying metabolism that contributes to the altered total C and N contents in mutant plants.

Our proposed energy conservation mechanism regulated by WRKY1 is also apparent in the regulation of a suite of dark inducible genes (DIN1, DIN4, DIN6/ASN1, and DIN10), in which WRKY1 represses these genes in the light. *At*DIN6/ASN1 in particular has been associated with both C and N signaling networks and energy conservation mechanisms in response to abiotic stress (Baena-Gonzales *et al*., 2007; Lam et al., 1998). The influence of WRKY1 on ASN1 under light and nitrogen stress is similar to the “low energy syndrome” (LES) described by (Tome *et al*., 2014 and Baena-Gonzalez and Sheen, 2008). The LES syndrome which plays a role in plant adaptation to stressful conditions in which non-specific stresses cause common energy deprivation responses. LES causes substantial perturbation of cellular processes including the arrest of metabolism and sugar storage and induction of catabolism, photosynthesis and remobilization of sugar (Tome *et al*., 2014; Baena-Gonzalez *et al*., 2008). *At*KIN10, an SNF1-related protein kinase, has been implicated as a factor controlling LES (Baena-Gonzalez et al 2007). Comparison of genes up and down regulated by *At*KIN10 (Baena-Gonzalez et al 2007) with genes up and down regulated in *wrky1* mutant plants revealed a unique and highly significant overlap (pval<0.001) between 81 genes up-regulated by *At*KIN10 and genes down-regulated by *WRKY1* (Supplemental Figure 7). GO term analysis of the 81 overlapping genes found a significant over-representation for the term “trehalose metabolic/biosynthetic processes” (pval=0.009). This is of particular interest since an association between trehalose metabolism and sugar-sensing in plants has recently been shown (Tsai and Gazzarrini, 2014), in which it is hypothesized that trehalose acts as a signal of sucrose availability (Schluepmann *et al*., 2003), and is shown to inhibit activity of the *At*SnRK1-KIN10 complex. Together, these results suggest that WRKY1 may play a role in mediating the LES syndrome in plants, having a potentially inverse but complementary role to the SnRK family of protein kinases. Although there were no observable differences in free Asn levels in two-week old mutant and WT plants, it is likely that there are differences in either protein-bound Asn levels or in free Asn levels but at a later stage since Asn is a known storage form of N (Lea et al., 2007; Gaufichon et al., 2016).

## CONCLUSION

The WRKY superfamily of transcription factors exist uniquely in plants, and are primarily associated with biotic and abiotic stress response (Rushton *et al*., 2010; Chen *et al*., 2013; Jia et al 2015). Our previous network analysis predicted that WRKY1 is a regulatory hub in the Arabidopsis N assimilation pathway, a component of primary metabolism. Here, we found that down-regulation of this single TF in a *wrky1-1* null mutant resulted in genome-wide transcriptional reprogramming of genes regulated by N and L signaling pathways, two essential plant response pathways. The phenotype of the wrky1 mutant shows that it plays a non-functionally redundant role compared to WRKY family members. Our assays show that *wrky1* mutants are affected in key metabolites of N-assimilation, including glutamine, aspartate, and glycine. Our results for carbon limitation suggest WRKY1 is involved in the low energy response pathways in Arabidopsis, and possibly other plant species. We speculate that WRKY1 is likely involved in mediating other abiotic stress response, as was recently shown by Qiao et al. (2016). Further study is required to investigate the full range of influence of WRKY1 on transcriptional regulation and resulting physiological phenotypes in response to environmental signals.

## MATERIALS AND METHODS

### Plant Material and Growth Conditions

*Arabidopsis thaliana* wild type (ecotype Columbia Col-0) seeds were obtained from Lehle Seeds, while *wrky1* T-DNA insertion lines were obtained from ABRC. Homozygous mutants were identified by PCR genotyping, using gene specific primers in combination with the T-DNA specific primer LBb1.3 (Supplemental Table 7). The lines SALK_016954 and SALK_136009 have a single polymorphism in the WRKY1 gene in the intron and promoter, respectively. However, the SALK_070989 line we used was recently shown by SALKSEQ to contain multiple polymorphisms. SALKSEQ_070989.0 and SALK_070989.56.00.x are T-DNA insertions in the intron sequence of *wrky1*, and both are present in our SALK_070989 line (Supplemental Figure 9). SALKSEQ_070989.1 is a T-DNA insertion in exon sequence of AT3G20460, a major facilitator superfamily protein. This insertion was not present in our SALK_070989 line (Supplemental Figure 9C), and the gene was not expressed based on microarray analysis. SALKSEQ_070989.2 is a T-DNA insertion in the intron of AT4G20300, serine/threonine-kinase, putative. This insertion was present in our SALK_070989 line (Supplemental Figure 9D). However, the expression of this gene only had a 1.2 fold change in expression compared to the WT and was not determined to be statistically significantly different between the WT and mutant line (Supplemental Table 2).

For steady-state or no-treatment experiments, wild type (Col-0) and homozygous mutant (SALK_070989; SALK_016954; SALK_136009) seeds were vapor-phase sterilized, vernalized for 3 days, then grown on basal MS media (Sigma M5524-1L), with 0.5 g/l MES hydrate (Sigma-Aldrich), 0.1% [w/v] sucrose, 1% [w/v] agar at pH 5.7. Plants were grown vertically on plates for 14 days in an Intellus environment controller (Percival Scientific, Perry, IA), under long-day (16 h light/8 h dark) conditions with light intensity of 50 μmol m-2s-1 at constant temp of 22□C. Seedlings were harvested two hours after start of light period and flash frozen in liquid N. For light treatments, Col-0 and SALK_070989 seedlings were grown exactly the same as no-treatment seedlings; however, at 13 DAP at start of the light period, half of the seedling plates were wrapped in a double layer of foil to extend darkness then placed back in the same chamber. On 14 DAP, seedlings were harvested two hours after start of the light period or putative start of light period for dark treated seedlings and immediately placed in liquid N. Dark treated seedlings were harvested at the same time as long day seedlings, but in complete darkness and flash frozen in liquid N. For nitrogen treatments, Col-0 and SALK_070989 seedlings were grown on basal MS media without N (custom GIBCO), supplemented with 1 mM KNO_3_^−^, 0.5 g/l MES hydrate (Sigma-Aldrich), 0.1% [w/v] sucrose, 1% [w/v] agar at pH 5.7. Seedlings were grown under the same conditions as the no-treatment seedlings for 14 days, then at start of the light period WT and mutant seedlings were transferred to either N-rich media (basal MS media without N (Phytotech), supplemented with 20 mM NH_4_NO_3_ plus 20 mM KNO_3_^−^, 0.5 g/l MES hydrate (Sigma-Aldrich), 0.1% [w/v] sucrose, 1% [w/v] agar at pH 5.7) or control media (basal MS media without N (Phytotech), supplemented with 20 mM KCl (molar equivalent for K in KNO_3_^−^), 0.5 g/l MES hydrate (Sigma-Aldrich), 0.1% [w/v] sucrose, 1% [w/v] agar at pH 5.7) for two hours then harvested and flash frozen in liquid N. For combined light and nitrogen treatments, half of the seedlings received extended dark treatment on 13 DAP as done for the light treatments, while the other half remained under normal light/dark regime. Nitrogen treatments were performed as before on 14 DAP in both light and dark conditions at start of light period. For all treatments, shoots and roots were harvested separately, and subsequent analyses were performed on shoot tissue only.

### RNA isolation, RT-qPCR, and Microarray

RNA from 3 biological replicates from each experiment was extracted from shoots using an RNeasy Mini Kit with RNase-free DNaseI Set (QIAGEN) and quantified on both a Nanodrop 1000 spectrophotometer (Thermo Scientific) and a Bioanalyzer RNA Nano Chip (Agilent Technologies). RNA was converted to cDNA (Thermoscript kit, Invitrogen) then analyzed by RT-qPCR using LightCycle FastStart DNA MasterPLUS SYBR Green I kit (Roche) with a LightCycler 480 (Roche, Mannheim, Germany). RT-qPCR primers are listed in Supplemental Table 7. Then, a 100 ng aliquot of total RNA was converted into cDNA, amplified and labeled with GeneChip 3’ IVT Express Kit Assay (Affymetrix). The labeled cDNA was hybridized, washed and stained on an ATH1-121501 Arabidopsis Genome Array (Affymetrix) using a Hybridization Control Kit (Affymetrix), a GeneChip Hybridization, Wash, and Stain Kit (Affymetrix), a GeneChip Fluidics Station 450 and a GeneChip Scanner (Affymetrix).

### Analysis and clustering of microarray data

Microarray intensities were normalized using the GCRMA (http://www.bioconductor.org/packages/2.11/bioc/html/gcrma.html) package in R (http://www.r-project.org/). For the steady-state experiment, differentially expressed genes for each mutant genotype were determined by Rank-Product (Breitling *et al*., 2004) and raw p-values were adjusted by False Discovery Rate (FDR) with a cutoff of 5%. For light-only and nitrogen-only experiments, differentially expressed genes were determined by two-way ANOVA with genotype and either light or nitrogen as factors. A gene was identified as differentially expressed if the FDR corrected p-value of ANOVA models was less than 0.01 and the p-value of the interaction coefficient, genotype and light or nitrogen (genotype × N or genotype × light), was less than 0.02. Tukey’s honest significant difference test (TukeyHSD) was used for multiple comparison to identify interaction term means that were significantly different from each other greater than the expected standard error. Only unambiguous probes were included. Multiple Experiment Viewer software (TIGR; http://www.tm4.org/mev/) was used to create heat maps and perform cluster analysis using Quality Threshold Clustering (QTC) with Pearson Correlation, HCL: average linkage method, and diameter 0.1. The significance of overlaps of gene sets were calculated using the GeneSect (R)script (15) using genes that are represented on the microarray as background. The significance of overrepresented Gene Ontology (GO) term analysis was performed with BioMaps (VirtualPlant 1.3) (Katari et al, 2010) using genes that are represented on the microarray as background in which the p-value of over-representation was measured by Fisher Exact Test with FDR correction and p-value cutoff of 0.01 or as otherwise indicated in figures. All microarray data have been deposited into GEO (http://www.ncbi.nlm.nih.gov/geo): GSE76278.

Three *wrky1* mutant T-DNA insertion lines were used to understand the core regulatory role of WRKY1 in response to nitrogen and light perturbations (see Plant Material and Growth Conditions). Microarray analysis was done for all three mutant lines plus wild type for “steady-state” and individual “light” and “nitrogen” treatments. Probes were normalized using the GCRMA method. Probes with more than one Present Call (P) in at least one group of replicates and a standard deviation greater than 0 were kept for further analysis. ANOVA with model simplification followed by Tukey HSD was performed for these three experiments in R using reshape2 (Wickham, 2007) and tidyverse (Wickham, 2017) packages and the GCRMA (Wu & Irizarry; MacDonald & Gentry, 2018), affy (Gautier et al., 2004), and BiocGenerics packages from Bioconductor. This analysis revealed that the three mutants lines respond in the same way to light and/or nitrogen perturbations, which is different from the WT response (Supplemental Figures 12 and 13), meaning that the same genes are either up or down-regulated across mutant genotypes in response to the treatment. Based on this analysis we were confident presenting results from the most severe *wrky1* mutant (*wrky1-1*) and WT control plants.

### Sequential ANOVA for combined experiment

For the combined N and light experiment, differentially expressed genes were determined by a three-way ANOVA with genotype, N, and light as factors. The ANOVA model was adjusted by FDR at cutoff of 1%, and genes significantly regulated by the interaction of genotype × light × nitrogen were selected with a p-value (ANOVA after FDR correction) cutoff of 0.01. Genes with significant three-way interaction were subjected to two-way ANOVA in which genotype × N interactions were explored across levels of the light variable, resulting in “Dark” and “Light” ANOVA models, similar to (threewayanova.htm. UCLA: Statistical Consulting Group; fromhttps://stats.idre.ucla.edu/spss/faq/how-can-i-explain-a-three-way-interaction-in-anova-2/, last accessed March 5, 2018). Genes with significant two-way interactions (ANOVA p-value <0.05 after model FDR correction, cutoff 5%) from Dark and Light ANOVA models were subjected to one-way ANOVA in which genotype factor was explored across levels of the N variable, resulting in “Dark Nitrogen”, “Dark Control”, “Light Nitrogen”, and “Light Control” ANOVA models. Genes with p-value < 0.05 (ANOVA after model FDR correction, cutoff 5%) were considered to have a significant genotype effect. Heat maps, cluster analysis, GO term analysis, and gene set overlap analysis were all performed as described above.

### Gene Network Analysis

Analysis of the nitrogen regulatory subnetwork was performed as described in Gutierrez *et al*., 2008, except that only one regulatory binding site was required for protein:DNA edges. The nitrogen × light crosstalk network was constructed from the 724 genes with a significant three-way interaction term for genotype × nitrogen × light. The Arabidopsis multi-network (VirtualPlant 1.3) was queried with this list of genes, and only significant correlation and protein:DNA regulatory edges were included in the network using Pearson correlation (cutoffs 1 to 0.7 or −1 to −0.7, with *P* ≤ 0.01). Networks were generated using the “Gene Networks” tool in the VirtualPlant system (www.virtualplant.org). Networks are visualized in Cytoscape 3.2.1.

### Promoter Analysis

The 2kb 5’ end upstream of the transcription start site (TSS) were considered the putative promoters regions of genes of interest. These regions were analyzed for known cis-regulatory element over-representation within a group of genes using Elefinder (http://stan.cropsci.uiuc.edu/cgi-bin/elefinder/compare.cgi), which returns an Expect value (e-value) that indicates how likely the result would be returned by chance based on the binomial distribution.

### Principle Component Analysis

Forty experiments (22 from LxNxG experiment, 12 from LxG experiment, and 6 from steady state) were re-normalized together using gcrma. The NxG hybridizations are the same as light treatments in the LxNxG. Normalized expression values were centered and used for principal component analysis using the prcomp function in R. The summary function in R was used to obtain the information regarding the % variance explained.

### Elemental Analysis

Arabidopsis thaliana lines Col-0 and SALK_070989 were grown on autoclaved soil (Sunshine Mix LC1) and fertilized with either ½ MS media minus C and N or ½ MS media supplemented with 50 mM NH_4_NO_3_. Total C, H, and N were determined by elemental analysis using an Exeter Analytical CHN Analyzer (Model CE440). Dried samples (30 mg FW, which is approximately 1.5 mg DW) were weighed in consumable tin capsules and purged with helium prior to combustion in pure oxygen under static conditions. Results were statistically analyzed using two-way analysis of variance.

### Metabolite Analysis

Arabidopsis thaliana lines Col-0 and SALK_070989 were grown on 50mL of Murashige and Skoog modified basal-salt mixture (Phytotech Labs M531) containing 1% w/v sucrose and either 0.5mM or 10mM NH4NO3 solution containing 20g/L BD bacto agar in a 100 × 100 × 15mm square petri dish with grid (Light Labs D210-16), three biological replicates each. Plants were grown for 14 days in Percival growth chamber (Percival Scientific) under long-day (16h light/8h dark) conditions with a light intensity of 120 μmolm-2s-1 and at a constant temperature of 22□C. Seedlings were harvested two hours after the start of the light period on the 14th day. The shoots from each plate were cut off, placed in an Eppendorf tube and flash frozen in liquid nitrogen. The samples were then stored at −80□C.

Metabolites were extracted based on the method outlined by Fiehn et al., 2008. The extraction solvent was prepared by mixing isopropanol/acetonitrile/water at the volume ratio of 3:3:2. For amino acid analysis, the concentrated samples were fractionated as outlined by Orlova et al., (2006). 1mL of water was added to each sample and vortexed until the residue was resuspended. 25μL of 10mM ribitol and 10mM alpha-aminobutyric acid was added as internal standards.

Samples were derivatized as outlined by Fiehn et al., 2008. An Agilent 7890B/7693 GC-MS system was used with a fused silica capillary column SPB-35 column (30 m × 320 μm × 0.25 μm; Supelco 24094). 1μL of each sample was injected using a splitless mode at 230□C. Helium (ultra high purity) was applied as the carrier gas using constant flow mode. The MS transfer line, ion source and quadrupole were kept at 250□C, 250□C and 150□C respectively. The GC oven was set to an initial temperature of 80□C and held for 2min. The temperature then increased at a rate of 5□C/min until a max temperature of 275□C and held for 6min. The MS was set to scan mode, and set to detect compounds eluting from 50-600m/z.

Agilent MassHunter Qualitative Analysis B.07.00 was used to obtain peak areas. Metabolite peaks were normalized to the internal standard and quantified as nmol/g FW. Statistical analysis (two-way ANOVA) was performed using R 3.5.2.

## Acknowledgements

The authors gratefully acknowledge Dr. Ying Li and Daniel Tranchina for advising on data analysis; and Dr. Gabriel Krouk for advice.

## Author contributions

AMC designed the research, performed the research, analyzed data, and wrote the paper. GC designed the research and wrote the paper. MK analyzed data and wrote the paper. RP assisted AMC with experiments. SH performed experiments and assisted AMC and MK to analyze data.

## FIGURE LEGENDS

**Supplemental Figure 1.** *WRKY1* regulatory subnetwork (VirtualPlant 1.1). Green lines indicate positive correlation and red lines indicate negative correlation of putative TF targets. Arrows indicate activation and flat lines indicate repression. Black lines indicate metabolic reactions. Correlation cutoff (≥0.7 or ≤ −0.7) and P-value ≤ 0.01.

**Supplemental Figure 2.** Relative expression of *WRKY1* in WT (Col-0) and *wrky1* T-DNA mutants measured by RT-qPCR; *wrky1-2* is SALK_016954; *wrky1-3* is SALK_136009; *wrky1-1* is SALK_070989. Error bars are standard error of the mean; three biological replicates.

**Supplemental Figure 3. Full cluster analysis of genes with significant Genotype × Light interaction. A.** Cluster analysis of genes with significant (pval<0.02, FDR 5%) Genotype × Light interaction effect (1567 genes). **B.** GO term analysis of gene clusters with significant GxL effect. *wrky1 = wrky1-1*; shaded areas = dark conditions.

**Supplemental Figure 4.** Crosstalk network of genes with significant three-way interaction (GxNxL). Nodes (580) are connected genes (transcription factor = triangle; metabolic = blue square; protein coding = purple square), edges (3110) are correlation based on co-expression (e-value cutoff = 0.01; green is negative; red is positive; increasing opacity corresponds to decreasing level of significance) and binding site over-representation, in which the target gene has at least one binding site for the transcription factor (Nero *et al*., 2009).

**Supplemental Figure 5.** Principal component analysis of all experiments identifies Light as the first principal component, explaining 49% of the variance, followed by Form of Nitrogen as the second principal component, explaining 30% of the variance. All experiments are labeled with L/D for Light or Dark, Col/WF for Wild type or *wrky1-1* Mutant, N/C for Nitrogen or Control, and finally with the replicate number. Steady state experiments are labeled with an upside down purple triangle, Light × Genotype experiments are labeled with red triangle, and finally the experiment with three factors (Light (L), Nitrogen (N), and Genotype (G)) are labeled with a blue diamond. Nitrogen × Genotype experiments is a subset of this dataset (Light only). It is clear from this figure that the first principal component is Light where all dark experiments are on the right and Light are on the left. And the second principal component is the form of Nitrogen treatment – Transient (bottom) and Constant (top).

**Supplemental Figure 6.** Representative genes of the WRKY1 mechanism in the dark, which are significant for the Light × Nitrogen × Genotype interaction shown in Figure 7. * indicates statistically significant difference at pval<0.01 as determined by sequential ANOVA (see Materials and Methods). D = Dark; N = Nitrogen treatment; C = KCl treatment; Mut = *wrky1-1*; WT = Col-0.

**Supplemental Figure 7.** Significance of overlaps (pval<0.001, number of overlapping genes inside parentheses) of *WRKY1* regulated (genes mis-regulated in *wrky1-1* mutant plants) and *At*KIN10 regulated (Baena-Gonzalez *et al*., 2007) gene sets, calculated using the GeneSect (R)script using the microarray as background. *At*KIN10 was overexpressed in protoplasts while *WRKY1* was down-regulated in whole plants. Therefore, the genes up-regulated by *At*KIN10 are down-regulated by *WRKY1*. Above the diagonal and in yellow; p-value <0.05, and the size of the intersection is higher than expected. Below the diagonal and in blue; p-value <0.05, and the size of the intersection is lower than expected (Katari *et al*., 2010).

**Supplemental Figure 8:** Measured metabolite levels (mean ± SD nmols/mg) in Col-0 and wrky1-1. **A.** asparagine; **B.** glutamate; **C.** fructose; **D.** glucose; **E.** threonic acid; **F.** serine; **G.** threonine; and **H.** glycine under nitrogen treatment of 0.5mM KNO^3−^ and 10.0mM KNO_3_^−^ (Tukey’s HSD, p-value < 0.05).

**Supplemental Figure 9: SALK_070989 genotyping gels.** 2% Agarose gel electrophoresis of PCR amplified products using the respective PCR primer set for each polymorphism. Lane L is a 1kB DNA size ladder for each image. **A.** SALK_070989.56.00.X. Lanes 1-7 are products using the forward and reverse primers, while lanes 8-14 are products using the LBb1.3 and reverse primers. Lanes 1 and 8 are blank PCR samples. Lanes 2-5 and 9-12 are Col-0 samples. Lanes 6-7 and 13-14 are SALK_070989 samples. **B.** SALKSEQ_070989.0 Lanes 1-7 are products using the forward and reverse primers. Lane 1 is a blank PCR sample. Lanes 2-5 are Col-0 samples. Lanes 6-7 are SALK_070989 samples. **C.** SALKSEQ_070989.0. Lanes 8-14 are products using the LBb1.3 and the reverse primers. Lane 8 is a blank PCR sample. Lanes 9-12 are Col-0 samples. Lanes 13-14 are SALK_070989 samples. **D.** SALKSEQ_070989.1. Lanes 1-4 are products using the forward and reverse primers, while lanes 5-8 are products using the LBb1.3 and reverse primers. Lanes 1 and 5 are blank PCR samples. Lanes 2-3 and 6-7 are Col-0 samples. Lanes 4 and 8 are SALK_070989 samples. **E.** SALKSEQ_070989.2. Lanes 1-7 are products using the forward and reverse primers, while lanes 8-14 are products using the LBb1.3 and reverse primers. Lanes 1 and 8 are blank PCR samples. Lanes 2-5 and 9-12 are Col-0 samples. Lanes 6-7 and 13-14 are SALK_070989 samples.

**Supplemental Figure 10: SALK_016954 and SALK_136009 genotyping gels.** 2% Agarose gel electrophoresis of PCR amplified products. **A.** SALK_016954. Lanes 1-4 are PCR amplified products using the forward and reverse primers, while lanes 5-8 are PCR amplified products using the LBb1.3 and reverse primers. Lanes 1 and 8 are blank PCR samples. Lanes 2 and 6 are Col-0 samples. Lanes 3-4 and 7-8 are SALK_016954 samples. Lane L is a 1kb DNA size ladder. **B.** SALK_136009. Lanes 1-4 are PCR amplified products using the forward and reverse primers, while lanes 5-8 are PCR amplified products using the LBb1.3 and reverse primers. Lanes 1 and 8 are blank PCR samples. Lanes 2 and 6 are Col-0 samples. Lanes 3-4 and 7-8 are SALK_136009 samples. Lane L is a 1kb DNA size ladder.

**Supplemental Figure 11:** A. QT clustering of differentially expressed genes under dark treatment (53% of D.E. genes). B. QT clustering of differentially expressed genes under light treatment (52% of D.E. genes). **1**. Col-0; **2**. *wrky1-2*; **3**. *wrky1-3*; **4**. *wrky1-1*

**Supplemental Figure 12:** A. QT clustering of differentially expressed genes under 20 mM NH_4_NO_3_, 20 mM KNO_3_ (85% of genes). B. QT clustering of differentially expressed genes under 20 mM KCl (67% of genes).

## TABLES

**Supplemental Table 1. Putative targets of WRKY1 identified from gene network analysis.** See: Supplemental Tables 1-6.xlsx

**Supplemental Table 2.** Significantly regulated genes from the steady-state (no treatment) conditions. Rank product pairwise comparisons between WT (Col-0) and wrky1 mutant lines (SALK_016954; SALK_136009; SALK_070989); genes have significant change in expression if pvalue<0.01, FDR cutoff = 5%. See: Supplemental Tables 1-6.xlsx

**Supplemental Table 3**. BioMaps output for 29 transcription factors in the crosstalk network.

**Supplemental Table 4.** Significantly regulated genes from the nitrogen treatment conditions; Control = 20 mM KCl; Treatment = (20 mM NH_4_NO_3_ + 20 mM KNO_3_^−^). Two-way ANOVA: WT (Col-0) and wrky1 mutant (SALK_070989); genes have significant change in expression if pvalue<0.02, ANOVA model FDR cutoff = 1%. See: Supplemental Tables 1-6.xlsx

**Supplemental Table 5.** Significantly regulated genes from the light treatment conditions; Control = long day conditions; Treatment = extended dark. Two-way ANOVA: WT (Col-0) and wrky1 mutant (SALK_070989); genes have significant change in expression if pvalue<0.02, ANOVA model FDR cutoff = 1%. See: Supplemental Tables 1-6.xlsx

**Supplemental Table 6.** Significantly regulated genes from the combined nitrogen and light treatment conditions; Controls = long day conditions; 20 mM KCl; Treatments = extended dark; (20 mM NH_4_NO_3_ + 20 mM KNO_3_^−^). Three-way ANOVA: WT (Col-0) and wrky1 mutant (SALK_070989); genes have significant change in expression if pvalue<0.01, ANOVA model FDR cutoff = 1%. See: Supplemental Tables 1-6.xlsx

**Supplemental Table 7.**
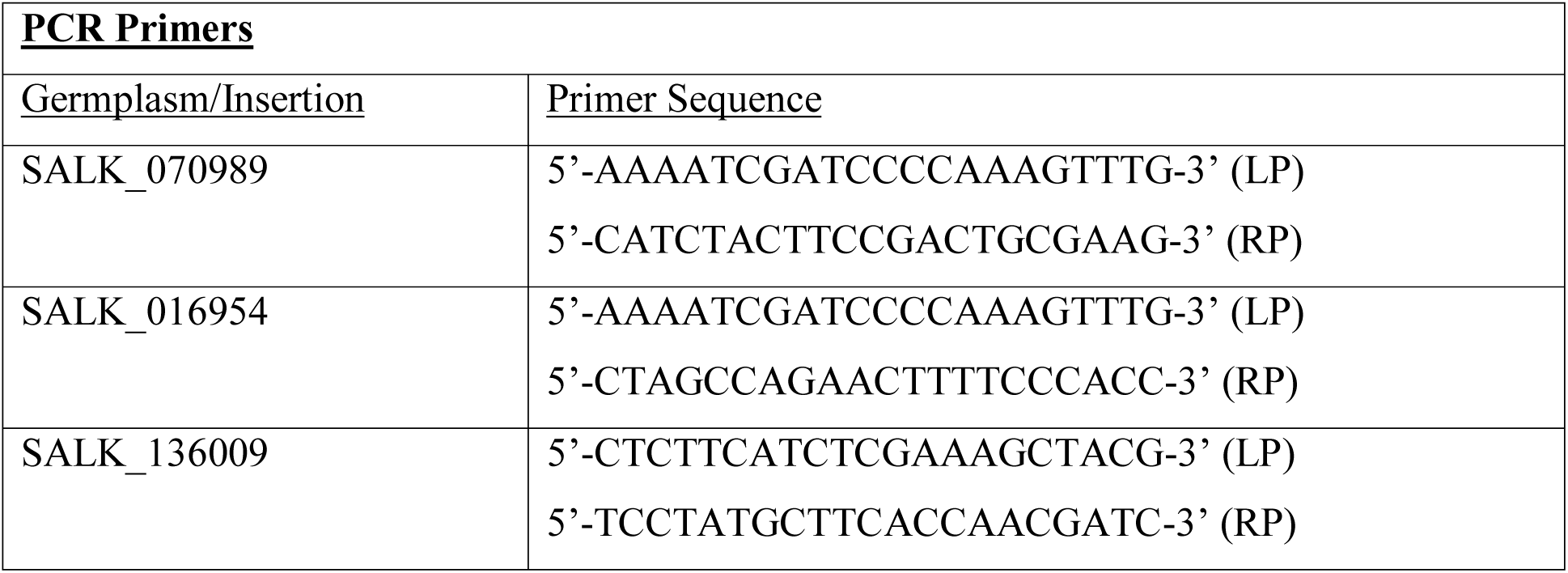
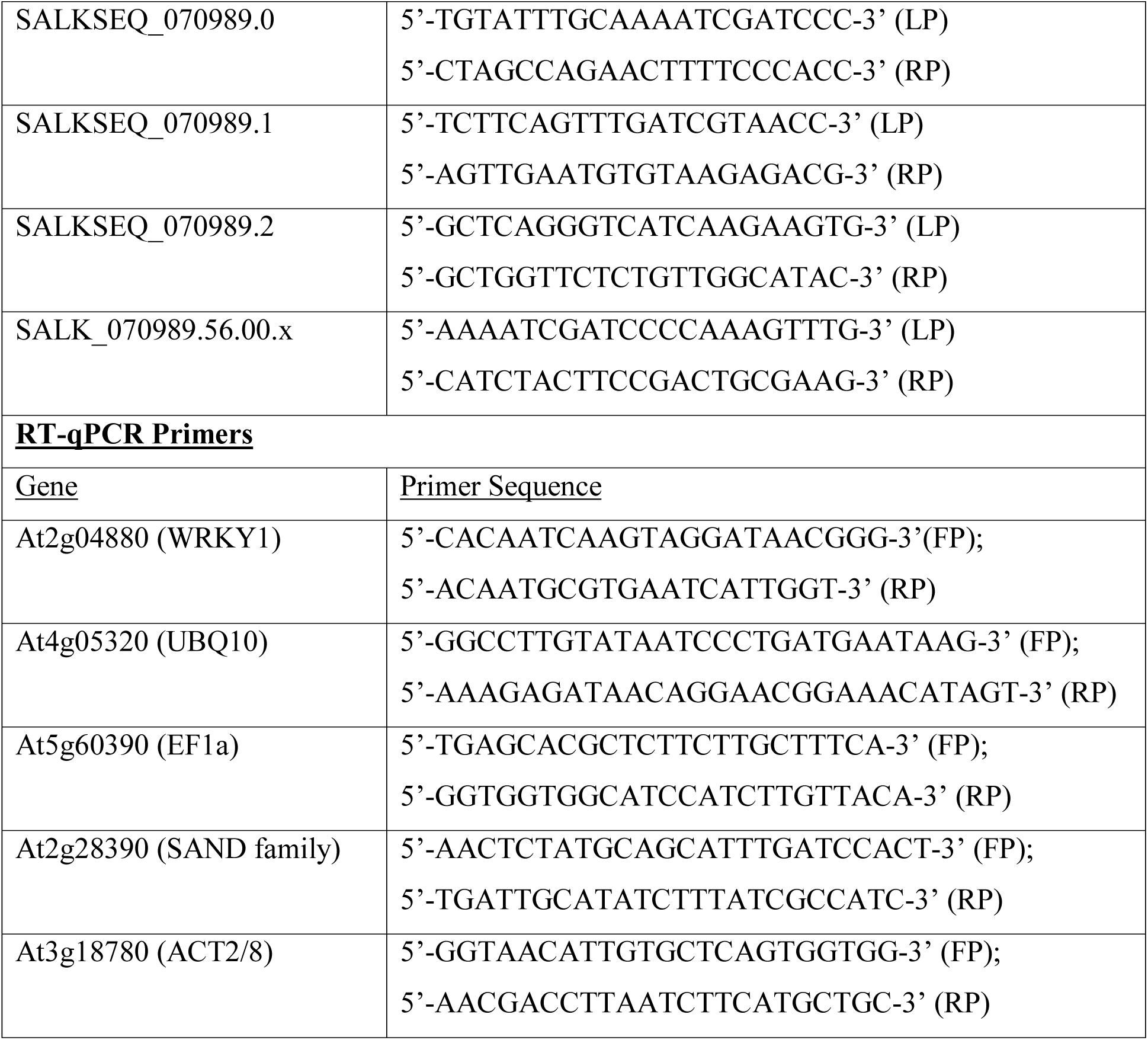
Primers used for genotyping germplasms and for RT-qPCR.

Supplemental Data Set 1. Relaxed network regulatory predictions from Gutierrez et al., 2008.

