## Supplemental Figure for "*WRKY1* mediates transcriptional crosstalk between light and nitrogen signaling pathways in *Arabidopsis thaliana*"

**Supplemental Figure 1.** WRKY1 regulatory subnetwork (VirtualPlant 1.1). Green lines indicate positive correlation and red lines indicate negative correlation of putative TF targets. Arrows indicate activation and flat lines indicate repression. Black lines indicate metabolic reactions. Correlation cutoff (≥0.7 or ≤ −0.7) and P-value ≤ 0.01.

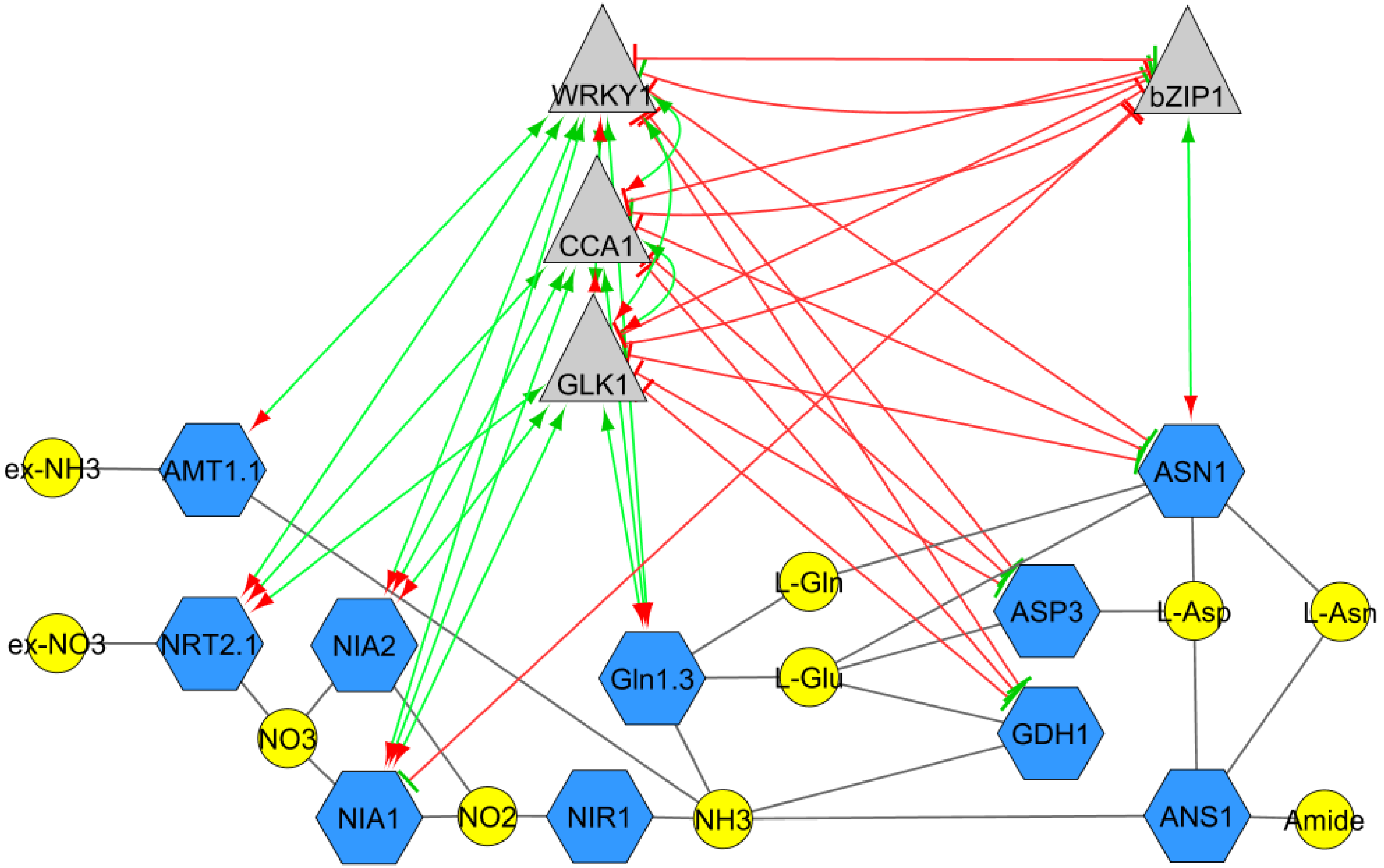

**Supplemental Figure 2.** Relative expression of WRKY1 in WT (Col-0) and *wrky1* T-DNA mutants measured by RT-qPCR. *wrky1-2* is SALK_016954; *wrky1-3* is SALK_136009; *wrky1-1* is SALK_070989. Error bars are standard error of the mean; three biological replicates.

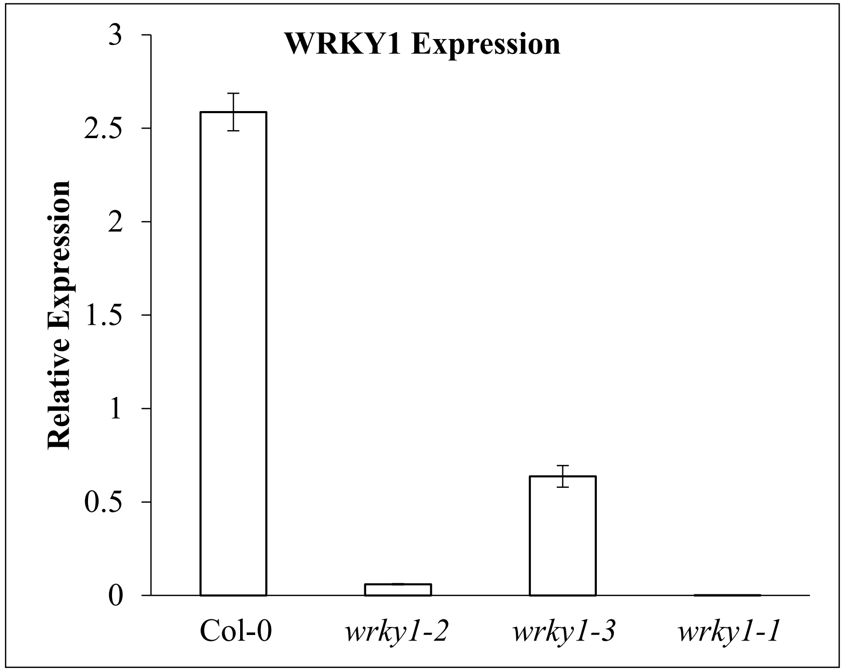

**Supplemental Figure 3. Full cluster analysis of genes with significant Genotype x Light interaction. A.** Cluster analysis of genes with significant (pval<0.02, FDR 5%) Genotype x Light interaction effect (1567 genes). **B.** GO term analysis of gene clusters with significant GxL effect. *wrky1* = *wrky1-1*; shaded areas = dark conditions.

**A**

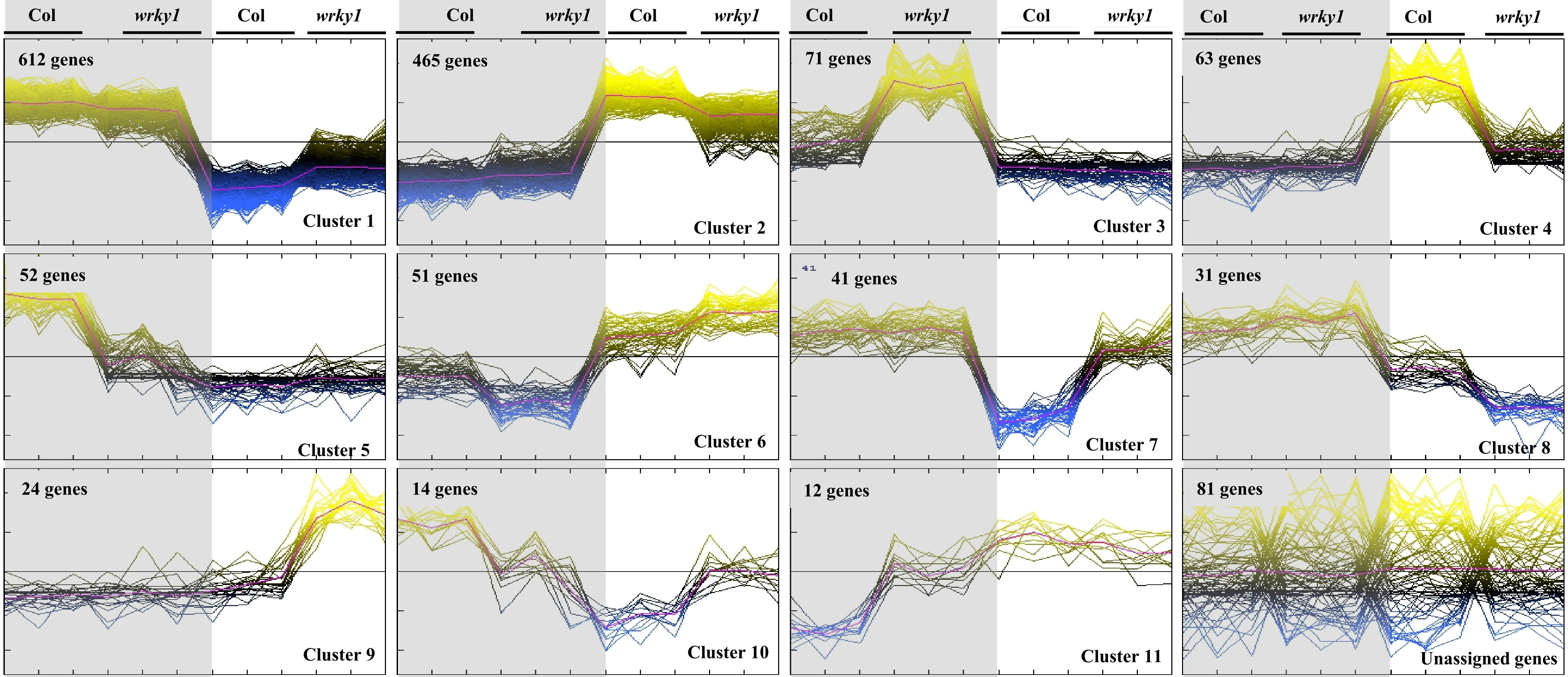


**B**

| Cluster | GO Term | pvalue | Cluster | GO Term | pvalue |
|---|---|---|---|---|---|
| 1 | Catabolic processes; response to stimuli | <0.01 | 7 | Regulation of cell morphogenesis | 0.05 |
| 2 | Metabolic processes; response to L; N compound metabolic processes | <0.01 | 8 | Coenzyme and pteridine-containing compound metabolic process | 0.07 |
| 3 | Secondary metabolic processes | 0.06 | 9 | Response to cold and to chemical stimulus | 0.07 |
| 4 | Trehalose, glycoside, and disaccharide biosynthetic processes | <0.01 | 10 | Biosynthetic processes: nucleotide, ATP, purine, ribonucleotide | <0.01 |
| 5 | Generation of precursor metabolites and energy; respiration; photosynthesis | <0.05 | 11 | Glycoside and carbohydrate metabolic processes | 0.04 |
| 6 | Response to endogenous, organic and hormone stimulus | <0.05 | | | |

**Supplemental Figure 4.** Crosstalk network of genes with significant three-way interaction (GxNxL). Nodes (580) are connected genes (transcription factor = triangle; metabolic = blue square; protein coding = purple square), edges (3110) are correlation based on co-expression (e-value cutoff = 0.01; green is negative; red is positive; increasing opacity corresponds to decreasing level of significance) and binding site over-representation, in which the target gene has at least one binding site for the transcription factor (Nero et al., 2009).

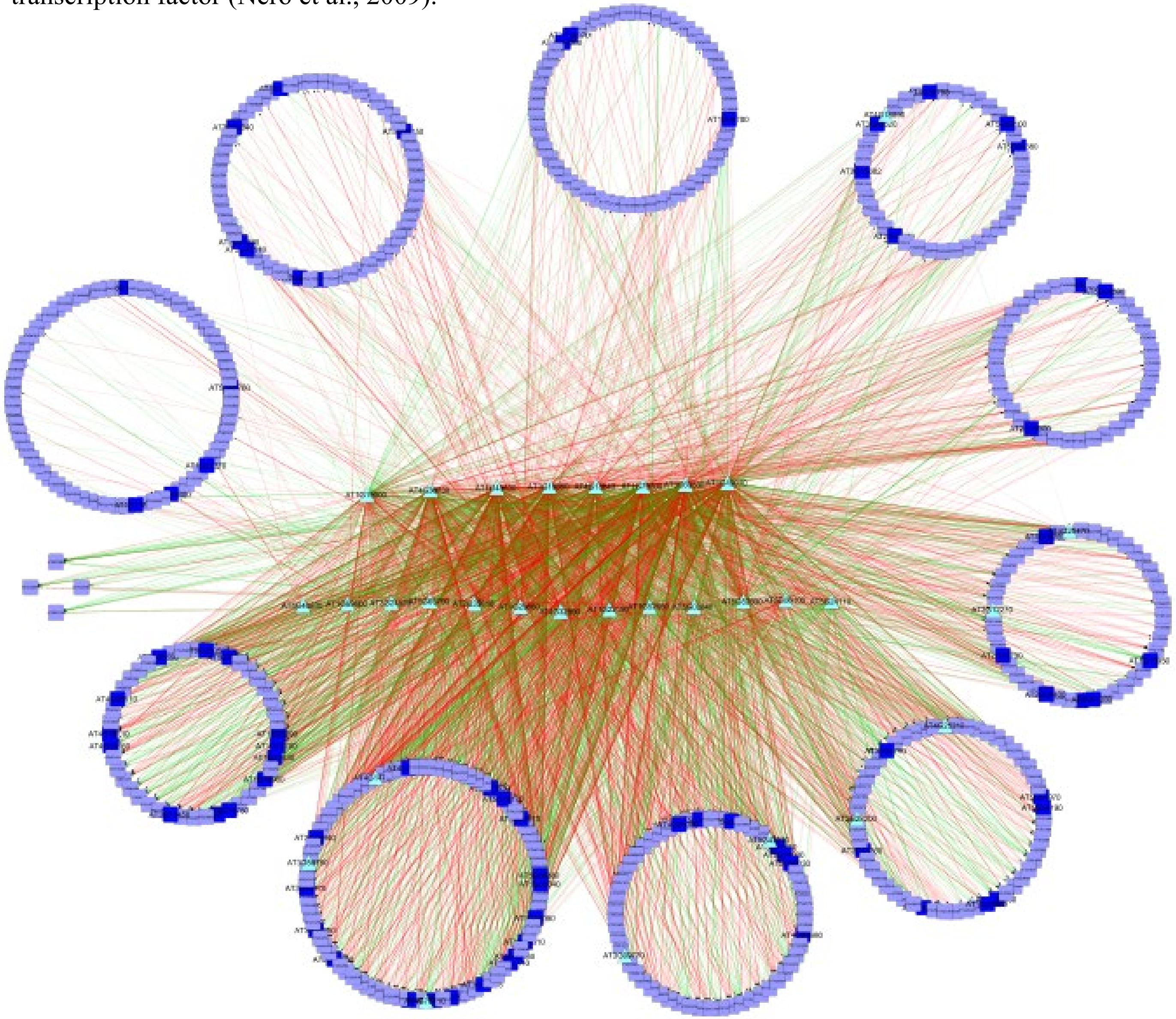

**Supplemental Figure 5.** Principal component analysis of all experiments identifies Light as the first principal component, explaining 49% of the variance, followed by Form of Nitrogen as the second principal component, explaining 30% of the variance. All experiments are labeled with L/D for Light or Dark, Col/WF for Wild type or *wrky1-1* Mutant, N/C for Nitrogen or Control, and finally with the replicate number. Steady state experiments are labeled with an upside down purple triangle, Light x Genotype experiments are labeled with red triangle, and finally the experiment with three factors (Light (L), Nitrogen (N), and Genotype (G)) are labeled with a blue diamond. Nitrogen x Genotype experiments is a subset of this dataset (Light only). It is clear from this figure that the first principal component is Light where all dark experiments are on the right and Light are on the left. And the second principal component is the form of Nitrogen treatment – Transient (bottom) and Constant (top).

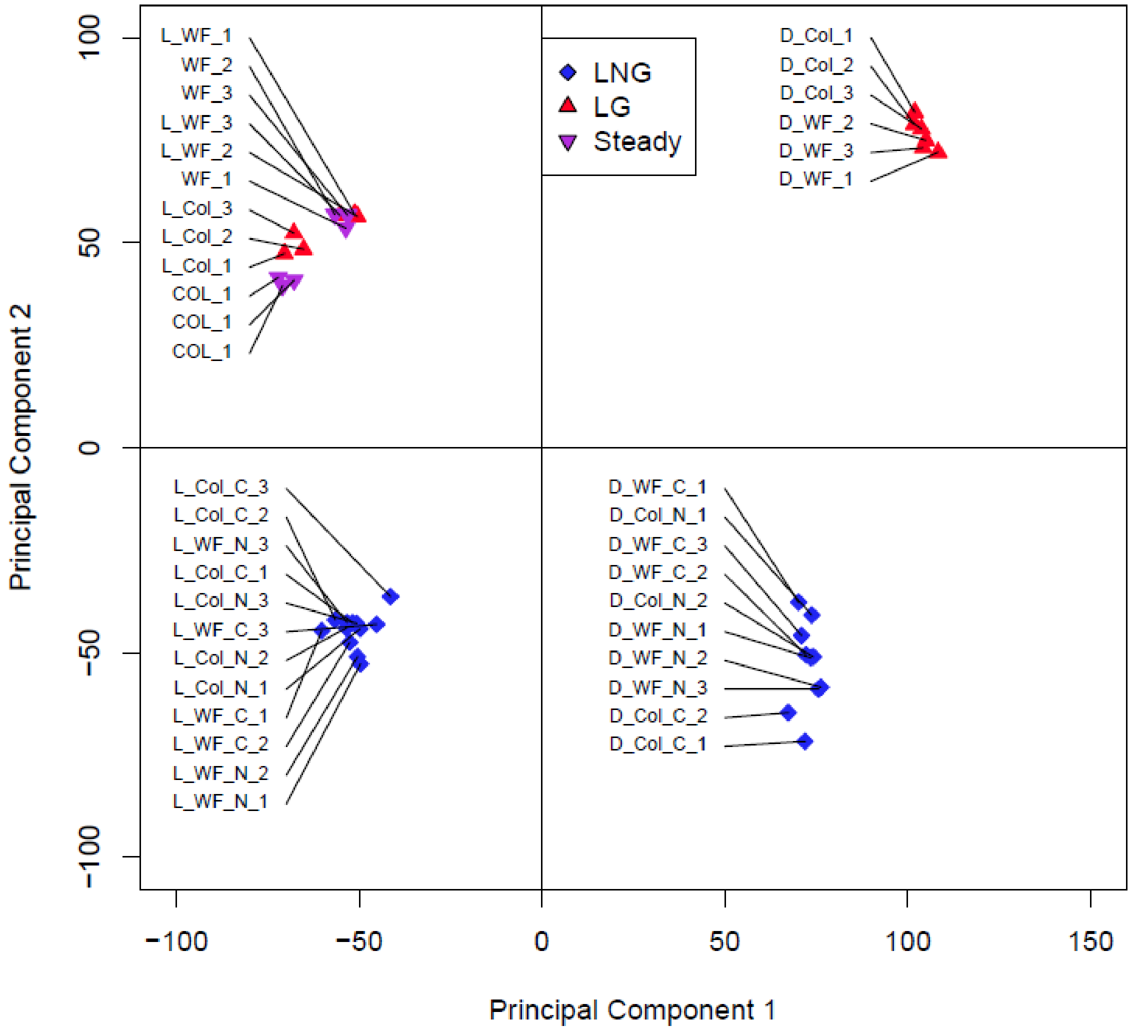

**Supplemental Figure 6.** Representative genes of the WRKY1 mechanism in the dark, which are significant for the Light x Nitrogen x Genotype interaction shown in Figure 7. * indicates statistically significant difference at pval<0.01 as determined by sequential ANOVA (see Materials and Methods). D = Dark; N = Nitrogen treatment; C = KCl treatment; Mut = *wrky1-1*; WT = Col-0.

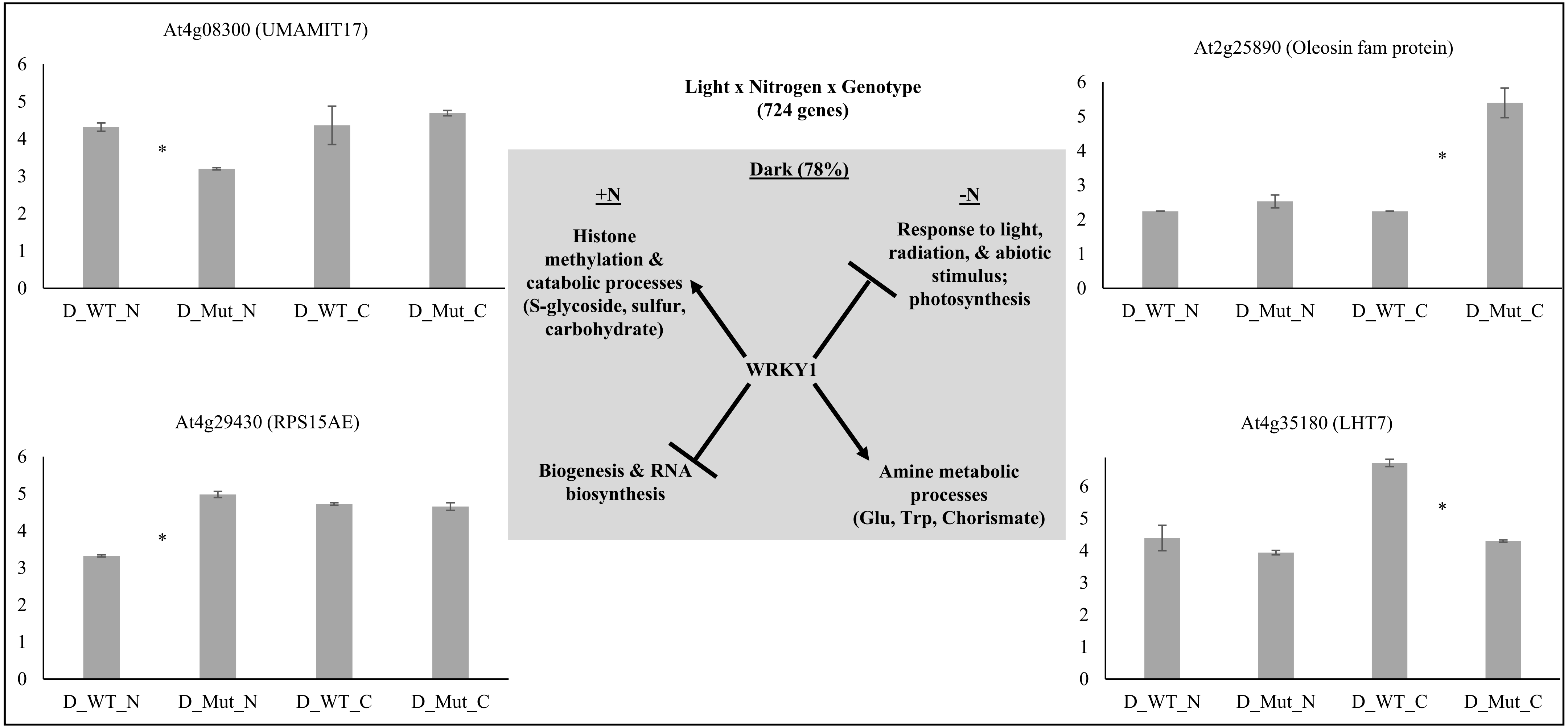

**Supplemental Figure 7.** Significance of overlaps (pval<0.001, number of overlapping genes inside parentheses) of WRKY1 regulated (genes misregulated in *wrky1-1* mutant plants) and KIN10 regulated (Baena-Gonzalez et al., 2007) gene sets, calculated using the GeneSect (R)script using the microarray as background. KIN10 was overexpressed in protoplasts while WRKY1 was down-regulated in whole plants. Therefore, the genes up-regulated by KIN10 are down-regulated by WRKY1. Above the diagonal and in yellow; p-value <0.05, and the size of the intersection is higher than expected. Below the diagonal and in blue; p-value <0.05, and the size of the intersection is lower than expected (Katari et al., 2010).

| | ***wrky1-1* Down** | ***wrky1-1* Up** | **KIN10 Down** | **KIN10 Up** |
|---|---|---|---|---|
| ***wrky1-1* Down** | | 1 (0) | 0.31 (3) | 0.61 (2) |
| ***wrky1-1* Up** | 0.39 (0) | | 0.99 (1) | <0.001 (81) |
| **KIN10 Down** | 0.86 (3) | 0.05 (1) | | 1 (0) |
| **KIN10 Up** | 0.66 (2) | 1 (81) | 0.001 (0) | |

**Supplemental Figure 8:** Measured metabolite levels (mean ± SD nmols/mg) in Col-0 and wrky1-1. **A.** asparagine; **B.** glutamate; **C.** fructose; **D.** glucose; **E.** threonic acid; **F.** serine; **G.** threonine; and **H.** glycine under nitrogen treatment of 0.5mM $KNO_3^-$ and 10.0mM $KNO_3^-$ (T-test, p-value: + < 0.2, ++ < 0.1, * < 0.05, ** < 0.01, *** < 0.001).

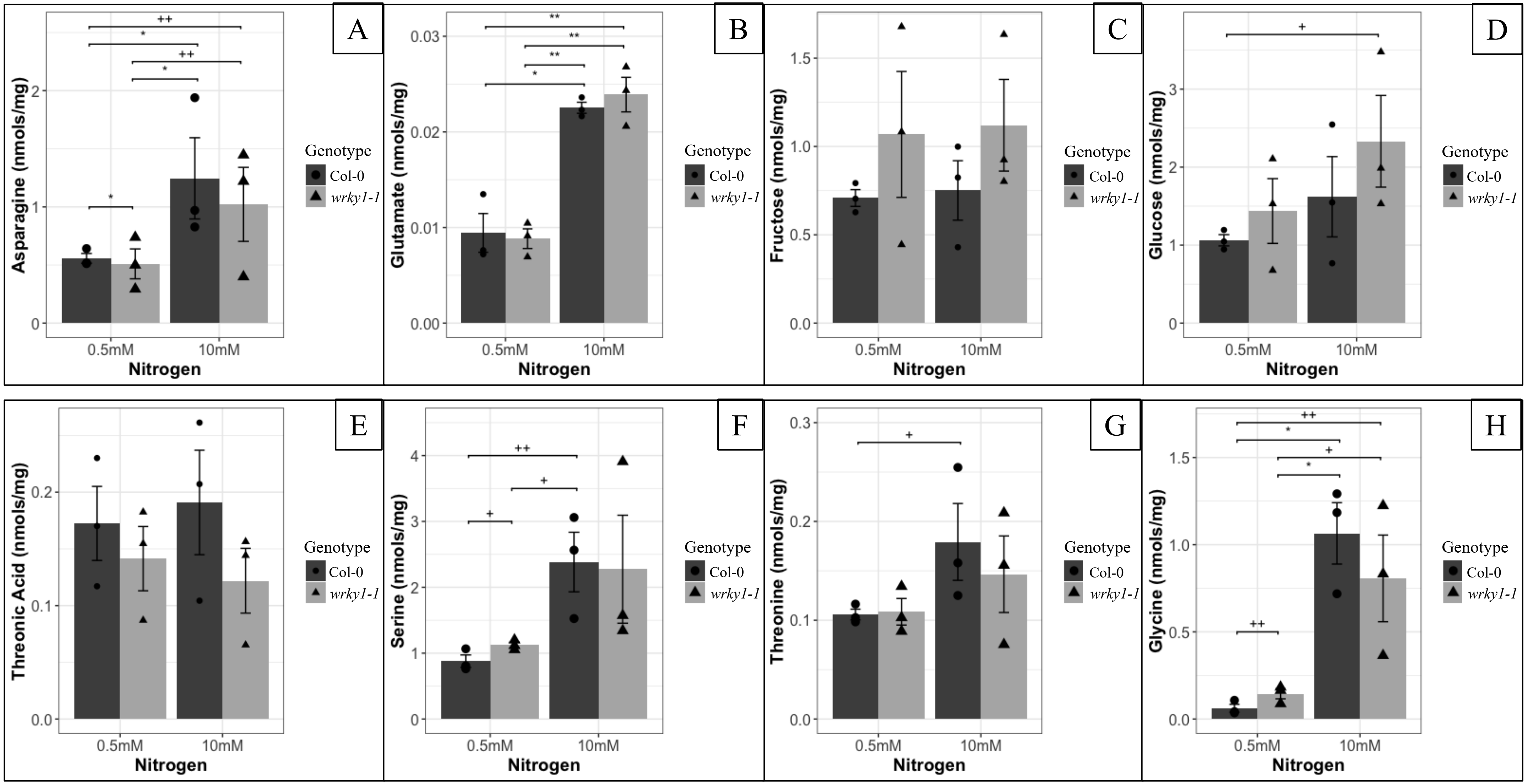

**Supplemental Figure 9**: **SALK_070989 genotyping gels**. 2% Agarose gel electrophoresis of PCR amplified products using the respective PCR primer set for each polymorphism. Lane L is a 1kB DNA size ladder for each image. **A.** SALK_070989.56.00.X. Lanes 1-7 are products using the forward and reverse primers, while lanes 8-14 are products using the LBb1.3 and reverse primers. Lanes 1 and 8 are blank PCR samples. Lanes 2-5 and 9-12 are Col-0 samples. Lanes 6-7 and 13-14 are SALK_070989 samples. **B.** SALKSEQ_070989.0 Lanes 1-7 are products using the forward and reverse primers. Lane 1 is a blank PCR sample. Lanes 2-5 are Col-0 samples. Lanes 6-7 are SALK_070989 samples. **C.** SALKSEQ_070989.0. Lanes 8-14 are products using the LBb1.3 and the reverse primers. Lane 8 is a blank PCR sample. Lanes 9-12 are Col-0 samples. Lanes 13-14 are SALK_070989 samples. **D.** SALKSEQ_070989.1. Lanes 1-4 are products using the forward and reverse primers, while lanes 5-8 are products using the LBb1.3 and reverse primers. Lanes 1 and 5 are blank PCR samples. Lanes 2-3 and 6-7 are Col-0 samples. Lanes 4 and 8 are SALK_070989 samples. **E.** SALKSEQ_070989.2. Lanes 1-7 are products using the forward and reverse primers, while lanes 8-14 are products using the LBb1.3 and reverse primers. Lanes 1 and 8 are blank PCR samples. Lanes 2-5 and 9-12 are Col-0 samples. Lanes 6-7 and 13-14 are SALK_070989 samples.

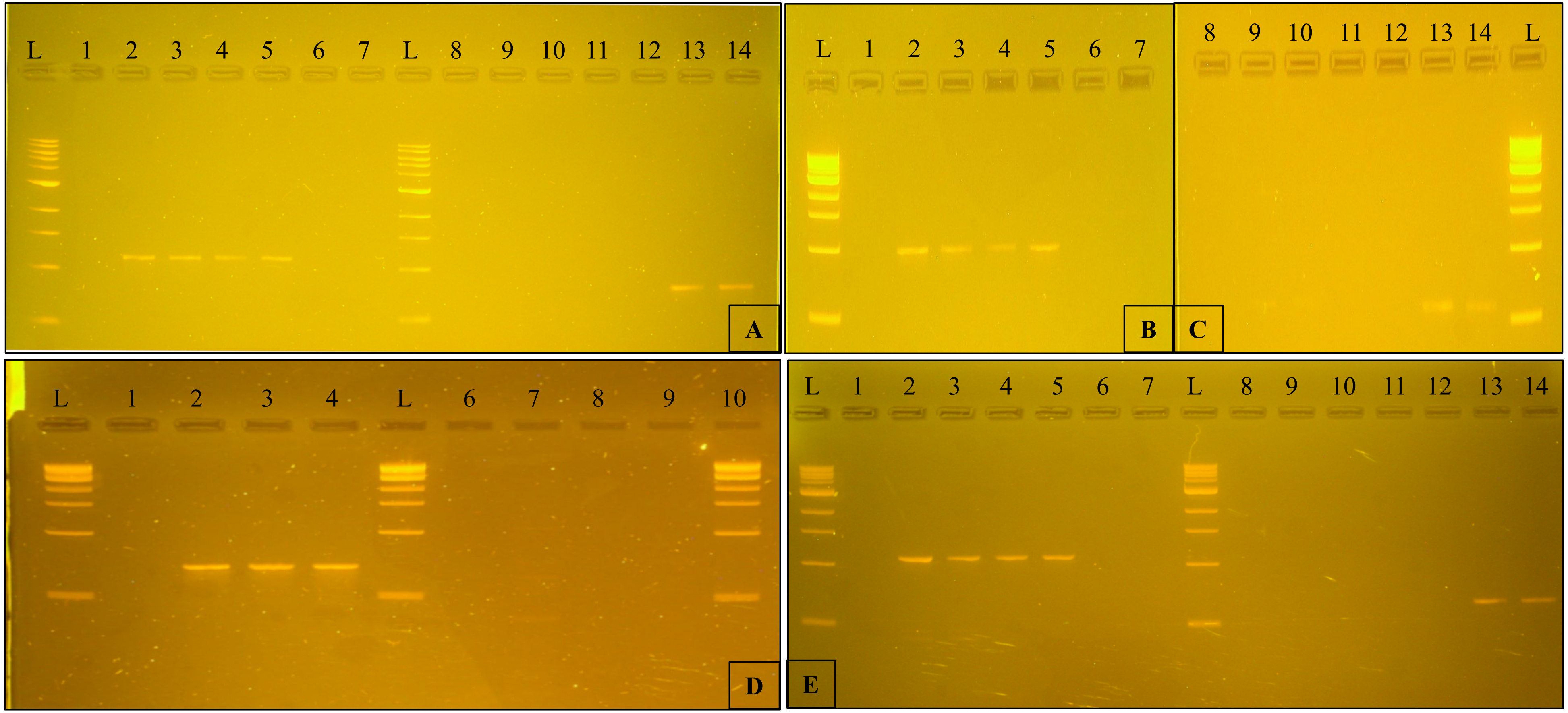

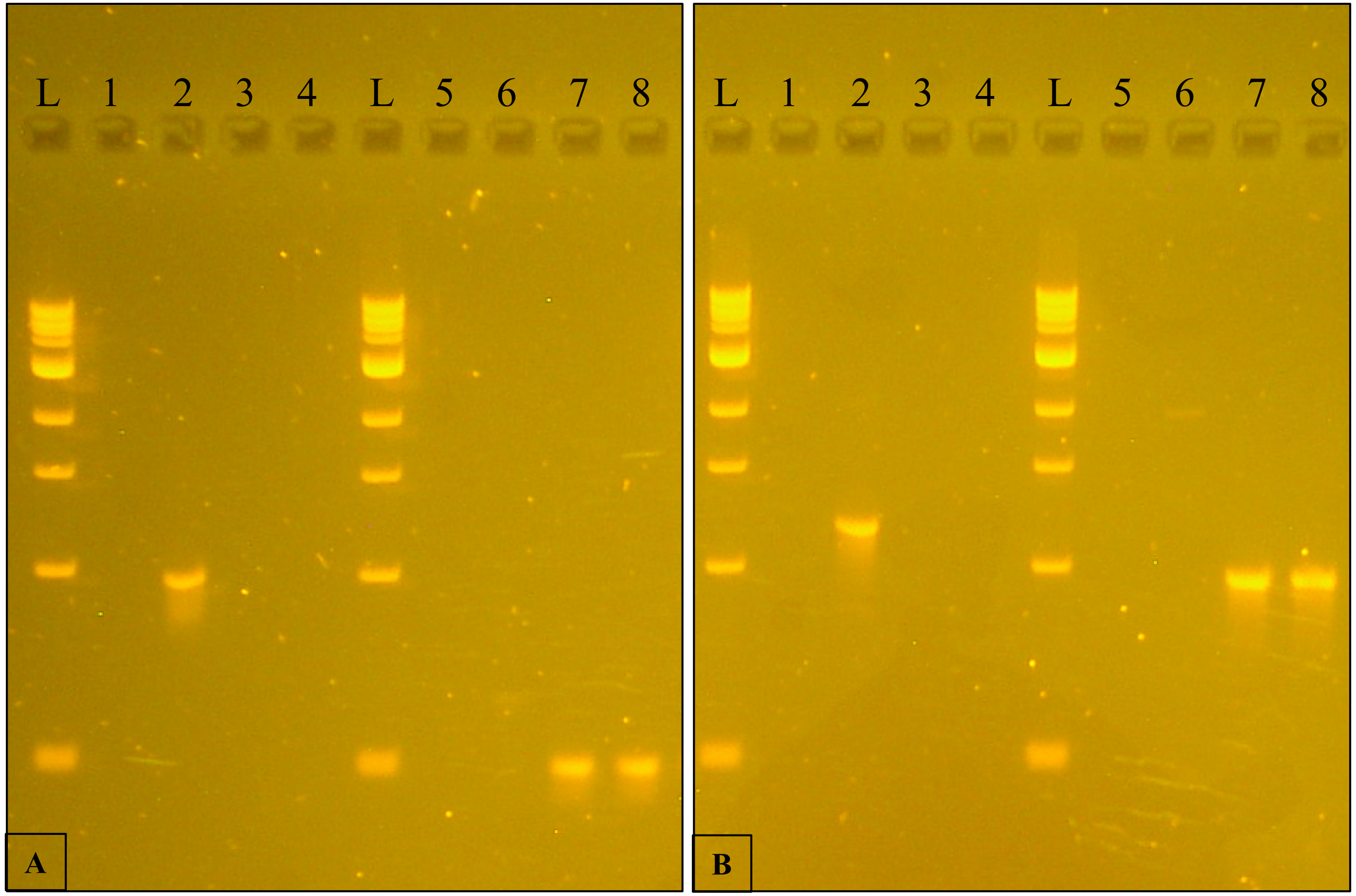


**Supplemental Figure 10: SALK_016954 and SALK_136009 genotyping gels.** 2% Agarose gel electrophoresis of PCR amplified products. **A.** SALK_016954. Lanes 1-4 are PCR amplified products using the forward and reverse primers, while lanes 5-8 are PCR amplified products using the LBb1.3 and reverse primers. Lanes 1 and 8 are blank PCR samples. Lanes 2 and 6 are Col-0 samples. Lanes 3-4 and 7-8 are SALK_016954 samples. Lane L is a 1kb DNA size ladder. **B.** SALK_136009. Lanes 1-4 are PCR amplified products using the forward and reverse primers, while lanes 5-8 are PCR amplified products using the LBb1.3 and reverse primers. Lanes 1 and 8 are blank PCR samples. Lanes 2 and 6 are Col-0 samples. Lanes 3-4 and 7-8 are SALK_136009 samples. Lane L is a 1kb DNA size ladder.

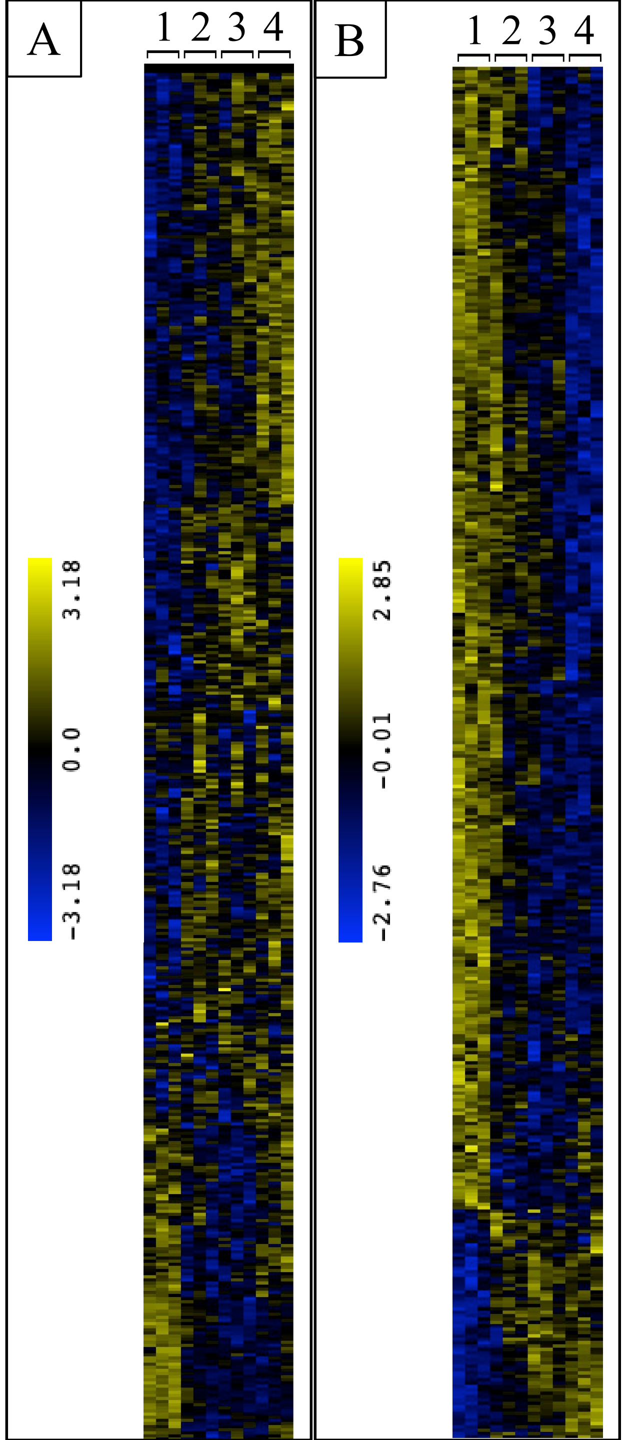


**Supplemental Figure 11: A.** QT clustering of differentially expressed genes under dark treatment (53% of D.E. genes). **B.** QT clustering of differentially expressed genes under light treatment (52% of D.E. genes). 1. Col-0; 2. *wrky1-2*; 3. *wrky1-3*; 4. *wrky1-1*

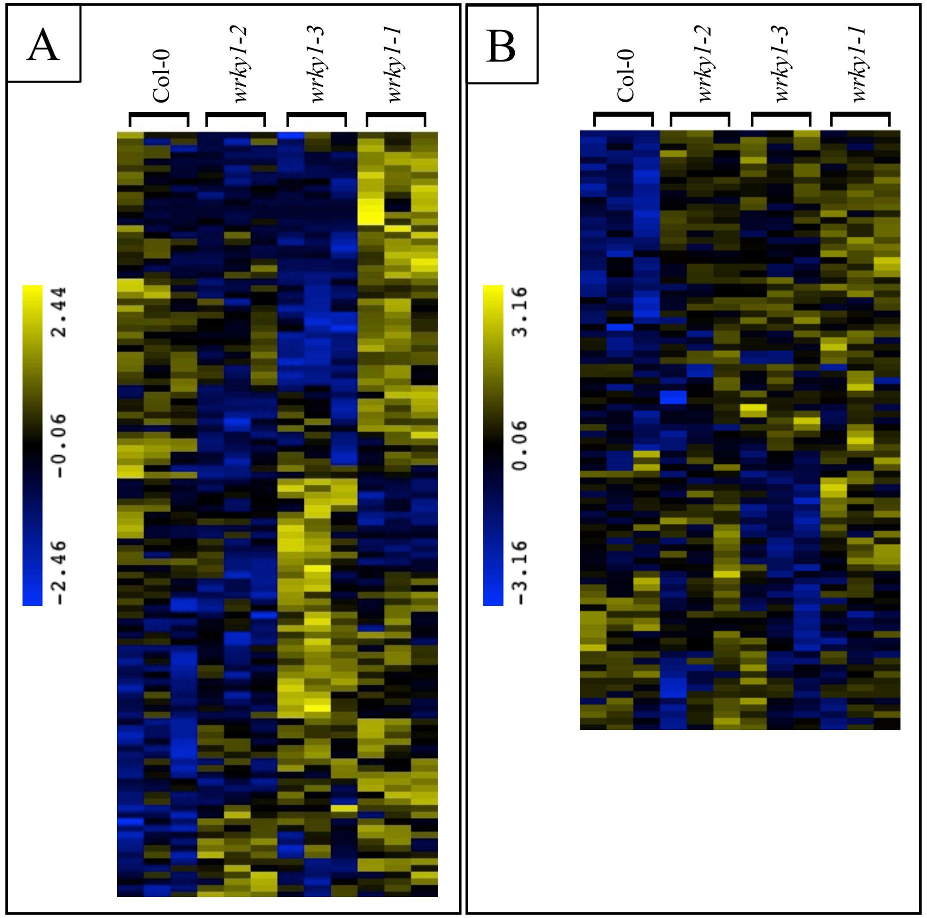


**Supplemental Figure 12**: **A.** QT clustering of differentially expressed genes under 20 mM $NH_4NO_3$, 20 mM $KNO_3$ (85% of genes). **B.** QT clustering of differentially expressed genes under 20 mM KCl (67% of genes).
